# The Vertebrate Codex Gene Breaking Protein Trap Library For Genomic Discovery and Disease Modeling Applications

**DOI:** 10.1101/630236

**Authors:** Noriko Ichino, MaKayla Serres, Rhianna Urban, Mark Urban, Kyle Schaefbauer, Lauren Greif, Gaurav K. Varshney, Kimberly J. Skuster, Melissa McNulty, Camden Daby, Ying Wang, Hsin-kai Liao, Suzan El-Rass, Yonghe Ding, Weibin Liu, Lisa A. Schimmenti, Sridhar Sivasubbu, Darius Balciunas, Matthias Hammerschmidt, Steven A. Farber, Xiao-Yan Wen, Xiaolei Xu, Maura McGrail, Jeffrey J. Essner, Shawn Burgess, Karl J. Clark, Stephen C. Ekker

## Abstract

The zebrafish is a powerful model to explore the molecular genetics and expression of the vertebrate genome. The gene break transposon (GBT) is a unique insertional mutagen that reports the expression of the tagged member of the proteome while generating Cre-revertible genetic alleles. This 1000+ locus collection represents novel codex expression data from the illuminated mRFP protein trap, with 36% and 87% of the cloned lines showcasing to our knowledge the first described expression of these genes at day 2 and day 4 of development, respectively. Analyses of 183 molecularly characterized loci indicate a rich mix of genes involved in diverse cellular processes from cell signaling to DNA repair. The mutagenicity of the GBT cassette is very high as assessed using both forward and reverse genetic approaches. Sampling over 150 lines for visible phenotypes after 5dpf shows a similar rate of discovery of embryonic phenotypes as ENU and retroviral mutagenesis. Furthermore, five cloned insertions were in loci with previously described phenotypes; embryos homozygous for each of the corresponding GBT alleles displayed strong loss of function phenotypes comparable to published mutants using other mutagenesis strategies (*ryr1b*, *fras1*, *tnnt2a, edar* and *hmcn1*). Using molecular assessment after positional cloning, to date nearly all alleles cause at least a 99+% knockdown of the tagged gene. Interestingly, over 35% of the cloned loci represent 68 mutants in zebrafish orthologs of human disease loci, including nervous, cardiovascular, endocrine, digestive, musculoskeletal, immune and integument systems. The GBT protein trapping system enabled the construction of a comprehensive protein codex including novel expression annotation, identifying new functional roles of the vertebrate genome and generating a diverse collection of potential models of human disease.

## Introduction

With the generation of more than 100 sequenced vertebrate genomes (Meadows & Lindblad-Toh, 2017), the current key question is how to determine the role(s) of uncharacterized gene products in specific biological and pathological processes. For example, genes associated with human disease are being discovered at a rapid rate. However, the biological functions underlying this linkage is often unclear (Kettleborough et al., 2013). Model system science using loss of function approaches has been essential to the annotation of the genome to date including the discovery of novel processes and the biological mechanisms underlying disease (Stoeger, Gerlach, Morimoto, & Nunes Amaral, 2018).

Among vertebrates, *Danio rerio* (zebrafish) has emerged as an outstanding model organism amenable to both forward and reverse genetic approaches. In addition, the natural transparency of the zebrafish embryo and larvae enables the unprecedented ability to non-invasively collect a rich set of expression data for the proteome and in the context of an entire living vertebrate. We describe here a 1000+ collection of zebrafish lines made using the Protein Trap Gene-Breaking Transposon (GBT;(Clark, Balciunas, et al., 2011)to develop such a codex for the comparative vertebrate genomics field (Meadows & Lindblad-Toh, 2017), (Clark, Balciunas, et al., 2011).

The initial pGBT-RP 2.1 (RP2.1) vector has several features that efficiency cooperate to report gene sequence, expression and function (Clark, Balciunas, et al., 2011). Two main reporter components include a 5’ protein trap and a 3’ exon trap, with the entire cassette flanked by inverted terminal repeats (ITR) of the mini*Tol2* transposon to effectively deliver the transgene as single copy integrations into the zebrafish genome. In cases where RP2 integrates in the sense orientation of a transcription unit, the protein trap’s splice acceptor overrides normal splicing of the transcription unit, creating a fusion between endogenous upstream exons and the monomeric RFP (mRFP) reporter sequences. The protein-trap domain in RP2.1 generates the expression profile, including subsequent protein localization and accumulation when a functional in-frame fusion between the start codon-deficient mRFP reporter and the tagged protein. Mutagenesis is accomplished by the strong internal polyadenylation and putative border element, effectively truncating the endogenously tagged protein. The GBT mutagenesis system represented the first step toward a ‘codex’ of protein expression and functional annotation of the vertebrate genome (Clark, Balciunas, et al., 2011).

We report here the development of a series of GBT protein trap vectors including versions to trap expression in each of the three potential reading frames. In addition, we modified the 3’ exon trap to use a localized BFP rather than the more commonly used GFP to more effectively use these lines in conjunction with other transgenic fish. We deployed these vectors at scale, generating over 1000 protein trap lines with visible mRFP expression at either 2dpf (end of embryogenesis) or 4dpf (larval stage), with 36% and 87% of the cloned lines showcasing to our knowledge the first described expression of these genes at these stages, respectively. We used forward and reverse genetic tests to assess the mutagenicity of these vectors, noting similar rates of visible phenotypes at 5dpf as ENU and retroviral screening tools. We re-isolated five previously described loci, and embryos homozygous for each of the corresponding GBT alleles displayed strong loss of function phenotypes comparable to these previously published mutants generated using other mutagenesis strategies (*ryr1b*, *fras1*, *tnnt2a, edar* and *hmcn1*). Molecular assessment after positional cloning shows that nearly all alleles cause at least a 99+% knockdown of the tagged gene. Interestingly, over 35% of the cloned loci represent 68 mutants in zebrafish orthologs of human disease loci, including nervous, cardiovascular, endocrine, digestive, musculoskeletal, immune and integument systems. The GBT protein trapping system enabled the construction of a comprehensive protein codex including novel expression annotation, identifying new functional roles of the vertebrate genome and generating a diverse collection of potential models of human disease.

## Materials and Methods

### Zebrafish husbandry

All zebrafish (*Danio rerio*) was maintained according to the guidelines and the standard procedures established by the Mayo Clinic Institutional Animal Care and Use Committee (Mayo IACUC). The Mayo IACUC approved all protocols involving live vertebrate animals (A23107, A21710 and A34513).

### Generating GBT constructs, RP2 and RP8 series

pGBT-RP8.2 and -RP8.3 were made by combining three restriction endonuclease fragments of pGBT-RP8.1, a 2.2 kb AflII to AgeI, a 0.7 kb EcoRI to SpeI, and a 3.0kb SpeI to AflII, with a short adapter to close the space between AgeI and EcoRI that effectively removed one or two thymine nucleotides just following the splice acceptor prior to the AUG-less mRFP cassette. For pGBT-RP8.2, Adapter-GBT(+2) was made by annealing oligos adapter-GBT(+2)-a [CCGGTTTTCTCATTCATTTACAGTCAGCCGG] and adapter-GBT (+2)-b [AATTCCGGCTGACTGTAAATGAATGAGAAAA]. For pGBT-RP8.3, Adapter-GBT(+3) was made by annealing oligos adapter-GBT (+3)-a [CCGGTTTTCTCATTCATTTACAGCAGCCGG] and adapter-GBT(+3)-b [AATTCCGGCTGCTGTAAATGAATGAGAAAA].

pGBT-RP2.2 and -RP2.3 were made by combining three restriction endonuclease fragments of pGBT-RP2.1 (Clark, Balciunas, et al., 2011), a 3.6kb BlpI to AgeI, a 1.9kb EcoRI to AvrII, and a 3.55kb AvrII to BlpI, with a short adapter to close the space between AgeI and EcoRI that effectively removed one or two thymine nucleotides just following the splice acceptor prior to the AUG-less mRFP cassette. For pGBT-RP2.2, Adapter-GBT(+2) was made by annealing oligos adapter-GBT(+2)-a and adapter-GBT (+2)-b. For pGBT-RP2.3, Adapter-GBT(+3) was made by annealing oligos adapter-GBT (+3)-a and adapter-GBT(+3)-b.

pGBT-RP8.1 was made by cloning a mini-intron derived from carp beta actin intron 1 into pGBT-RP7.1. The 234bp SalI to XhoI mini-intron fragment was isolated from pCR4-bactmIntron following digestion. The pGBT-RP7.1 plasmid was digested with XhoI so that the SalI to XhoI fragment was cloned between the gamma-crystallin promoter and nls tagBFP.

pCR4-bactmIntron was made by removing a 1.1kb internal portion of the carp beta actin intron 1 by digestion of pCR4-bact_I1 with BstBI and BssHII, followed by filling in 5’ overhangs and ligating remaining vector fragment.

pCR4-bact_I1 was cloning a PCR product containing the carp beta-actin intron into pCR4-TOPO (Invitrogen). The intron was amplified from pGBT-RP2.1 (Clark, Balciunas, et al., 2011) using MISC-bact_exon-F1 [CAGCTAGTGCGGAATATCATCTGCC] and MISC-bact_intron-R1 [CTTCTCGAGGTGAATTCCGGCTGAACTGTA] primers.

pGBT-RP7.1 was made by replacing a 501bp PstI to PstI fragment of pGBT-RP6.1 with a 480bp PstI to PstI fragment of pRP2.1. This changed the nucleotide sequence between the carp beta-actin splice acceptor to replicate the sequences in pGBT-RP2.1. pGBT-RP7.1 was never directly tested in zebrafish.

pGBT-RP6.1 was made by flipping the internal trap cassette relative the Tol2 inverted terminal repeats in pGBT-RP5.1. To do this, pGBT-RP5.1 was cut with EcoRV and SmaI. The 2.27kb EcoRV to SmaI vector backbone fragment, which included the ITRs, was ligated to the 3.51kb EcoRV to SmaI trap fragment. pGBT-RP6.1 was then selected based on the right ITR of Tol2 being in front of the RFP trap, which is the same orientation of pGBT-RP2.1.

pGBT-RP5.1 was made by cloning a PCR product with the AUG-less mRFP into pre(−1)GBT-RP5.1. The 698bp mRFP* PCR product was obtained by amplification of pGBT-R15 (Clark, Balciunas, et al., 2011) with CDS-mRFP*-F1 [AAGAATTCGAAGGTGCCTCCTCCGAGGATGTCATCAAGG] and CDS-mRFP-R1 [AAACTAGTCTTAGGCTCCGGTGGAGTGGCGG]. Prior to cloning the PCR mRFP* product was digested with EcoRI and SpeI to prepare the ends for subcloning into pre(−1)GBT-RP5.1 that was opened between the carp beta actin splice acceptor and the ocean pout terminator.

pre(−1)GBT-RP5.1 was made by cloning 1.2kb SpeI to AvrII fragment from pGBT-PX (Sivasubbu et al., 2006) that contained the ocean pout terminator into the SpeI site of pre(−2)GBT-RP5.1. The resulting products were screened for the proper orientation of the ocean pout terminator relative to the carp beta actin splice acceptor.

pre(−2)GBT-RP5.1 was made by inserting an expression cassette to make a 3’ poly(A) trap that makes blue lenses. A 1.15kb SpeI to BglII fragment from pKTol2gC-nlsTagBFP was cloned into pre(−3)GBT-RP5.1 that had been cut with AvrII and BglII. This moved the *Xenopus* gamma crystallin promoter driving a nuclear-localized TagBFP in front of the carp beta actin splice donor within pre(−3)GBT-RP5.1 to create a localized BFP poly(A) trap signal replacing the ubiquitous GFP signal that was in pGBT-RP2.1.

pre(−3)GBT-RP5.1 was made by cloning a 492bp XmaI to NheI scaffold fragment from pUC57-I-SceI_loxP_splice into pKTol2-SE (Clark, Balciunas, et al., 2011) opened with XmaI and NheI. pUC57-I-SceI_LoxP_Splice contains a synthetic sequence (see below) cloned into pUC57 (Genscript). The scaffold contains an I-SceI site; loxP site; carp beta actin splice acceptor; cloning sites for mRFP, ocean pout terminator, and BFP lens cassettes; carp beta actin splice donor; loxP site; and an I-SceI site. [cccgggatagggataacagggtaatataacttcgtatagcatacattatacgaagttatcgttaccacccactagcggtcagactgcagattgcagcacgaaacaggaagctgactccacatggtcacatgctcactgaagtgttgacttccctgacagctgtgcactttctaaaccggttttctcattcatttacagttcagcctgttacctgcactcaccgacaagctgttaccctggaattcgtttaaacactagtcaccggcgttcctaggttataagatctacctaaggtgagttgatctttaagctttttacattttcagctcgcatatatcaattcgaacgtttaattagaatgtttaaataaagctagattaaatgattaggctcagttaccggtcttttttttctcatttacactgagctcaagacgtctgataacttcgtatagcatacattatacgaagttattaccctgttatccctatggctagc]

### Generating GBT collection

Generation of the GBT collection was based on the prior described protocols (Clark, Balciunas, et al., 2011; Clark, Urban, Skuster, & Ekker, 2011; J. Ni et al., 2016).

### Fluorescent microscopy of mRFP reporter protein expression

Larvae were treated with 0.2 mM phenylthiocarbamide at 1 dpf to inhibit pigment formation. The anesthetized fish were mounted in 1.5% agarose (Fisher Scientific BP1360) prepared with 0.017mg/ml tricaine solution in an agarose column in the imaging chamber. The protocol of ApoTome microscopy was described in previous publication. (Clark, Balciunas, et al., 2011) For Lightsheet microscopy, larval zebrafish were anesthetized with 0.017g/ml tricaine (Ethyl 3-aminobenzoate methanesulfonate salt) in embryo water during imaging procedure. To capture RFP expression patterns of 2 dpf and 4 dpf larval zebrafish, LP 560 nm filter as excitation and LP 585nm as emission was used for Lightsheet microscopy.

The sagittal-, dorsal-, and ventral-oriented z-stacks of the mRFP expression were captured at either 50x magnification using an ApoTome microscope (Zeiss) with a 5x/0.25 NA dry objective (Zeiss) or 50x magnification using a Lightsheet Z.1 microscope (Zeiss) 5x/0.16 NA dry objective. Each set of images were obtained from the same larva and the images shown are composites of the maximum image projections of the z-stacks obtained from each direction.

### Sperm Cryopreservation

Sperm collection and cryopreservation was initially based on the protocol described in (Draper & Moens, 2009) and moved to the ZIRC protocol described in (Matthews et al., 2018).

### Genomic DNA isolation

Genomic DNA was isolated from F1 fish tail biopsies to conduct next generation sequencing and from both WT and heterozygous larva to manually perform the PCR-based mRFP linkage analysis. Zebrafish larva were individually placed to 0.2 ml PCR tubes and sacrificed to extract genomic DNA in 50 mM NaOH for 20 min at 95 C^o^.

### PCR-based linkage analysis of GBT insertions loci

TALE-PCR: The protocol used was designed to amplify and clone junction fragments from Tol2-based gene-break transposons (GBT) in the zebrafish genome. Although modified, it is based on a protocols received from Alexi Parnov, Vladimir Korsch, and Karuna Sampath. The following primer mixtures (containing 0.4 μM GBT specific primer and 2 µM DP primer) were prepared: for primary PCR: 5R-mRFP-P1/DP1, 5R-mRFP-P1/DP2, 5R-mRFP-P1/DP3, 5R-mRFP-P1/DP4, 3R-GM2-P1/DP1, 3R-GM2-P1/DP2, 3R-GM2-P1/DP3, 3R-GM2-P1/DP4, 3R-tagBFP-P1/DP1, 3R-tagBFP-P1/DP2, 3R-tagBFP-P1/DP3, 3R-tagBFP-P1/DP4; for secondary PCR: 5R-mRFP-P2/DP1, 5R-mRFP-P2/DP2, 5R-mRFP-P2/DP3, 5R-mRFP-P2/DP4, 3R-GM2-P2/DP1, 3R-GM2-P2/DP2, 3R-GM2-P2/DP3, 3R-GM2-P2/DP4, 3R-tagBFP-P2/DP1, 3R-tagBFP-P2/DP2, 3R-tagBFP-P2/DP3, 3R-tagBFP-P2/DP4; for tertiary PCR: TAIL-bA-SA/DP1, TAIL-bA-SA/DP2, TAIL-bA-SA/DP3, TAIL-bA-SA/DP4, Tol2-ITR(L)-O1/DP1, Tol2-ITR(L)-O1/DP2, Tol2-ITR(L)-O1/DP3, Tol2-ITR(L)-O1/DP4, Tol2-ITR(L)-O3/DP1, Tol2-ITR(L)-O3/DP2, Tol2-ITR(L)-O3/DP3, Tol2-ITR(L)-O3/DP4. A total of 1 μl of primer mixtures were added to PCR reaction (total volume 25 µl). Cycle settings were as follows. Primary: (1) 95°C, 3 min; (2) 95°C, 20 sec; (3) 61°C, 30 sec; (4) 70°C, 3 min; (5) go to “cycle 2” 5 times; (6) 95°C, 20 sec; (7) 25°C, 3 min; (8) ramping 0.3°/sec to 70°C; (9) 70°C, 3 min; (10) 95°C, 20 sec; (11) 61°C, 30 sec; (12) 70°C, 3 min; (13) 95°C, 20 sec; (14) 61°C, 30 sec; (15) 70°C, 3 min; (16) 95°C, 20 sec; (17) 44°C, 1 min; (18) 70°C, 3 min; (19) go to “cycle 10” 15 times; (20) 70°C, 5 min; Soak at 12 °C. A total of 5 μl of the primary reaction was diluted with 95 µl of 10mM Tris-Cl or TE buffers and 1μl of the mixture was added to the secondary reaction. Secondary: (1) 95°C, 2 min (2) 95°C, 20 sec; (3)61°C, 30 sec; (4) 70°C, 3 min; (5) 95°C, 20 sec; (6) 61°C, 30 sec; (7) 70°C, 3 min; (8) 95°C, 20 sec; (9) 44°C, 1 min; (10) ramping 1.5°/sec to 70°C; (11) 70°C, 3 min; (12) go to “cycle 2” 15times; (13) 70°C, 5 min; Soak at 12°C. A total of 5 μl of the primary reaction was diluted with 95 µl of 10mM Tris-Cl or TE buffers and 1μl of the mixture was added to the tertiary reaction. Tertiary: (1) 95°C, 2 min; (2) 95°C, 20 sec; (3) 44°C, 1 min; (3) ramping 1.5°/sec to 70°C; (4) 70°C, 3 min; (5) go to “cycle 2” 32 times; (6) 70°C, 5 min; Soak at 12°C. Products of the secondary and tertiary reactions were separated by using 1-1.5% agarose gel. The individual bands from the “band shift” pairs were cut from the gel and purified by using QIAquick Gel Extraction Kit (QIAGEN, Germany), and sequenced by using ABI Cycle Sequencing chemistry (PE Applied Biosystems, CA) and an ABI Prism 310 Genetic Analyzer with Data Collection Software (PE Applied Biosystems, Foster City, CA) supplied by the producer.

5’ and 3’ RACE PCRs: The protocol used was designed to use cDNA to amplify and clone junction fragments of Tol2-based gene-break transposons (GBT) in the zebrafish genome. 5’ RACE allows PCR amplification of unknown sequence at the 5’ end of a cDNA as long as there is enough known sequence within the cDNA to design two antisense primers. Although modified, it is based on the protocol in described in (Clark, Balciunas, et al., 2011). The following primer mixtures were prepared: for primary PCR: 0.20 μM GBT specific primer (5R-mRFP-P1), and a mix of universal 5’ RACE primers 2.5 μM 5R-UP-S and 0.5 μM 5R-UP-L. Secondary reaction: 25 μM GSP (5R-mRFP-P2), and 25 μM universal primer 5R-N1. The reaction mix used was as follows: Primary: (25 μl reaction) Template (RR-cDNA) 2 μl, Bioline 5X myTaq buffer 5 μl, myTaq 0.25 μl, GSP 0.5 μl, Universal 5’ RACE primer mix (URS mix) 2.0 μl, Water 15.25 μl. Secondary: (50 μl reaction) Template (2:100 dilution 1° PCR Reaction), Bioline 5X myTaq buffer 10 μl, myTaq 0.3 μl, GSP-P2 0.9 μl, URS-P2 0.9 μl, Water 35.9 μl. Cycle Settings are as follows. Primary: (1) 95°, 3’; (2) 95°, 30“; (3) 65°, 30” -0.5°/cycle; (4) 70°, 2’; Go To (2) x 15 cycles; (5) 95°, 30”; (6) 57°, 30”; (7) 70°, 2’; Go To (5) x 20 cycles, (8) 70°, 10’; (9) Soak at 12°. Dilute 2µL of the primary PCR reaction with 198µL of 10mM Tris-Cl; 1mM EDTA pH8.0. Secondary: (1) 95° for 3’ (2) 95°, 30”; (3) 63°, 30” -0.5°/cycle; (4) 70°, 2’; Go To (2) x 10 cycles; (5) 95°, 30”; (6) 58°, 30”; (7) 70°, 2’; Go To (5) x 25 cycles; (8) 70°, 10’; Soak at 12°. After the PCR reactions finish 20µL of each sample were run on a 1.2% agarose gel. You may also run 10µL of undiluted primary PCR reactions on the same gel; however, most of the time the bands of interest do not all appear until after the nested PCR reactions have been run. The individual bands from the gel were excised and purified by using QIAquick Gel Extraction Kit (QIAGEN, Germany), and sequenced by using ABI Cycle Sequencing chemistry (PE Applied Biosystems, CA) and an ABI Prism 310 Genetic Analyzer with Data Collection Software (PE Applied Biosystems, Foster City, CA) supplied by the producer.

### Forward genetic screening with next-generation sequencing

Isolated genomic DNA (300-500ng) was digested with MseI, and BfaI in parallel for 3h at 37°C and heat inactivated for 10 min at 80°C. The digested samples from each enzyme were pooled with prealiquoted barcoded linker in individual wells. The T4 DNA ligase (New England Biolabs, Inc.) was added, and the reaction mix was incubated for 2 h at 16°C. The linker-mediated PCR was performed in two steps. In the first step, PCR was done with one primer specific to the 3’-ITR (5’-GACTTGTGGTCTCGCTGTTCCTTGG-3’) and the other primer specific to linker sequences (5’-GTAATACGACTCACTATAGGGC-3’) using the following conditions: 2 min at 95°C, 25 cycles of 15 sec at 95°C, 30 seconds at 55°C and 30 seconds at 72°C. The PCR products were diluted to 1:50 in dH2O, and a second round of PCR was performed using ITR (5’-TCACTTGAGTAAAATTTTTGAGTACTTTTTACACCTC-3’) and linker specific (5’ -GCGTGGTCGACTGCGCAT-3’) nested primers to increase sensitivity and avoid non-specific amplification using the following conditions: 2 min at 95°C, 20 cycles of 15 sec at 95°C, 30 seconds at 58°C and 30 seconds at 72°C. The nested PCR products from each 96-well plate are pooled and processed for Illumina library preparation as per manufacturer’s instructions.

### Protein classification of the cloned zebrafish genes

The 183 cloned zebrafish genes are classified by using PANTHER (Mi, Muruganujan, & Thomas, 2013). PANTHER provided protein classes of the molecule coded by the cloned zebrafish genes.

### Annotating human orthologues of GBT-tagged genes and disease-causing genes

The human orthologues of 192 cloned zebrafish genes were mainly collected by using a data mining tool, ZebrafishMine (Van Slyke et al., 2018) supported by the ZFIN database. In some cases, the candidates of human orthologues unlisted in ZFIN database were manually searched by using both Ensembl (https://useast.ensembl.org/index.html) and InParanoid8 (http://inparanoid.sbc.su.se/cgi-bin/index.cgi) databases. In parallel, the candidates were manually identified by the result of BLASTP assembled with human proteins and by the result of an online synteny analysis tool, SynFind (https://genomevolution.org/CoGe/SynFind.pl). If the candidate multiply hit in those manual assessments, it was annotated as a human orthologue. The human phenotype data caused by mutations of 68 human orthologues were collected by using another data mining tool, BioMart (http://useast.ensembl.org/biomart/martview/cfe15ead83199a0b7c7997f5a4ce9e6b) supported by Ensembl database.

### Finding Disease Models in Vertebrates

Mouse models were found by using both descriptions of animal models in Online Mendelian Inheritance in Man (OMIM; https://www.omim.org/) and in Mouse Genome Informatics (MGI; http://www.informatics.jax.org/humanDisease.shtml). MGI provided the details of mouse models of human disease, such as the number of models have been established. Zebrafish model were also found by using both OMIM and the Zebrafish Information Network (ZFIN; https://zfin.org/). ZFIN provided all data of fish strains listed in this database.

### Gene expression profiling of the cloned zebrafish genes

The cloned genes with unpublished expression data were isolated by using “Gene Expression” tool of ZebrafishMine (Van Slyke et al., 2018). In parallel, some published expression data were also manually searched from ZFIN database or in some references. To isolate the gene with the expression localized in the tissues or the organs shows abnormalities in the causing diseases of the human orthologues, the mRFP reporter expression patterns of the cloned genes 2 and 4 dpf are manually analyzed using zfishbook database (Clark, Argue, Petzold, & Ekker, 2012).

## Results

### The features of GBT constructs RP2 and RP8 – capturing all three proteomic reading frames

In our previous study, we reported the intronic-based gene-breaking transposons (GBTs) as effective and revertible loss-of function tools for zebrafish (Clark, Balciunas, et al., 2011). The main features of the RP2.1 vector system are as follows (Fig1A): 1) Genetically engineered cargo is flanked by miniTol2 sequences necessary and sufficient for Tol2 transposase-mediated transposition, an efficient transgenesis vector in zebrafish (Kawakami et al., 2004); (Balciunas et al., 2006); (Urasaki, Morvan, & Kawakami, 2006); 2) protein trap that enables *in vivo* expression selection of the vertebrate proteome. The addition of the AUG-free mRFP reporter has yielded an effective protein trap for both organ-specific and subcellular localization of the tagged locus (Petzold et al., 2009); (Clark, Balciunas, et al., 2011); (Liao et al., 2012);(Xu et al., 2012);(Ding et al., 2013); (Westcot et al., 2015); 3) Mutagenic transcriptional terminator. The 5’ cassette is a combination of a strong splice-acceptor (SA), poly adenylation signal (pA), and putative border element (red octagon) in conjunction with a start codon (AUG)-free monomeric red fluorescent protein (mRFP) reporter. These elements have been shown to be effective as a transcriptional stop in zebrafish by hijacking endogenous splicing (Sivasubbu et al., 2006). These elements are very effective at inducing a quantitative knockdown in all 26 lines assessed to date using qRT-PCR; 97% or higher knockdown in all lines (Clark, Balciunas, et al., 2011); (Ding et al., 2013); (Ding et al., 2016), this manuscript). The GBT mutagenesis system is thus an effective first step to creating a gene codex simultaneously combining expression and loss of function genetics. However, some limitations were noted with the initial RP2.1 vector – notably the effective trapping of transcripts without detectable expression of the mRFP reporter. Molecular cloning of these GFP+/RFP-lines demonstrated the fidelity of expression requiring the capture of an appropriate reading frame. RP2.1 was designed to use one main reading frame, and some lines with expression were noted to include the use of a secondary splice acceptor (data not shown). To maximize genome coverage of this insertional mutagen, we created all three reading frames of the RP2 and RP8 vector series (Fig. 1). 4) 3’ exon trap. These vectors also encodes a 3’ exon trap with preferential expression following intragenic insertions in zebrafish (Sivasubbu et al., 2006); (Petzold et al., 2009); (Clark, Balciunas, et al., 2011). The function of this cassette complements the obligate, in-frame protein trapping effect and is used for both quality control during mutagenesis and for genotyping of more weakly expressing protein trap alleles (Fig.1A). In the RP2 vector series, the nearly ubiquitous b-actin promoter drives expression of GFP. Expression of integrated GFP becomes detectable between early developmental stages such as seven- to eight-somite-stage and provides bright expression with a good signal-to-noise ratio at 25 hpf (Davidson AE et al., 2003). However, the ubiquitous GFP expression from the 3’ exon trap cassettes could interfere with another fluorescent marker system based on GFP labeling in further studies. The RP8 vector series includes all reading frames for the AUG-free mRFP reporter and a new 3’ exon trap cassette with expression of tagBFP driven by the lens-specific gamma-crystalline promoter (Fig. 1B). Using the tissue-specific reporter system with BFP is helpful to easily detect F1 founder with weak mRFP expression and to avoid interference with GFP-based multi-labeling purposes when crossed with other GFP-labeled transgenic fish lines. 5) Revertible mutagenic cassette. The flanking loxP sites enable reversion of the tagged locus by Cre – mediated recombination. This facilitates both somatic (Clark, Balciunas, et al., 2011); (Ding et al., 2013) and germline approaches (Petzold et al., 2009). We generated more than eleven hundred independent lines by using all six constructs of the GBT system (Supplemental Table 1). We conducted an initial screening expression of the mRFP fusion protein and showed that RP2 and RP8 vector series with all reading frames of mRFP reporter protein readily detects the distribution of the fusion proteins expressed from their own promoter in zebrafish (Supplemental Figure 1).

**Figure 1.**
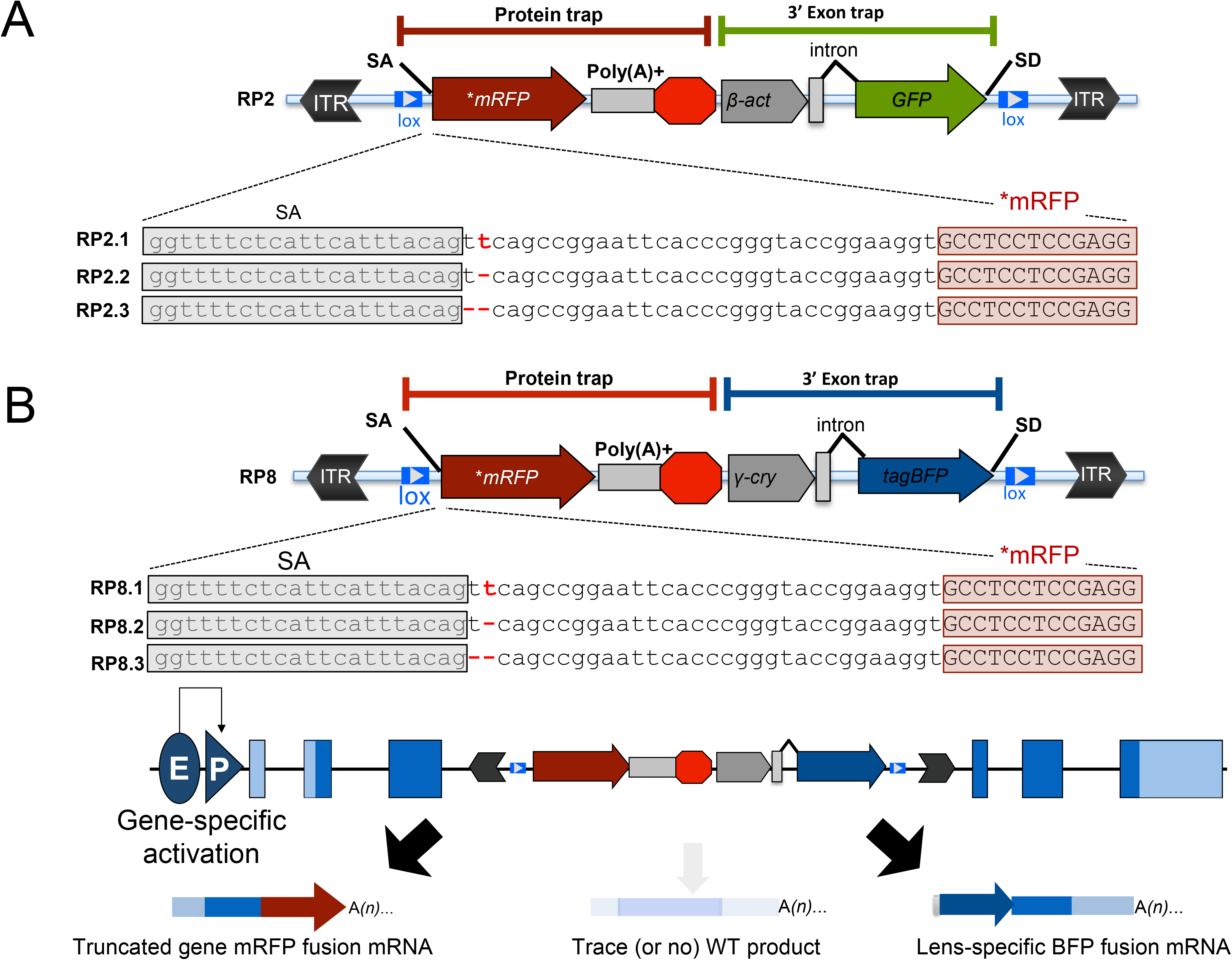
Schematic of the RP2 and RP8 gene-break transposon system with all three reading frames of AUG-less mRFP reporter. A. Schematic of the RP2 system fused with 3 reading frames of AUG-less mRFP reporter (RP2.1, RP2.2 and RP2.3). B. Schematic of the RP2 system fused with 3 reading frames of AUG-less mRFP reporter (RP8.1, RP8.2 and RP8.3). ITR, inverted terminal repeat; SA, loxP; Cre recombinase recognition sequence, splice acceptor; *mRFP’ AUG-less mRFP sequence; poly (A)+, polyadenylation signal; red octagon, extra transcriptional terminator and putative border element; *β-act*, carp beta-actin enhancer, SD, splice donor; E, enhancer; P, promoter; and WT, wild type.

### Annotation of protein localization and trafficking of the GBT strain collection

The ability to non-invasively obtain temporal and spatial expression pattern information is a key feature of these protein trap strains. Pilot data from our first RFP lines rapidly demonstrated this step was going to be a major bottleneck for our pipeline if we used standard documentation methods. Consequently, we established a capillary-based confinement and imaging protocol (SCORE imaging; (Petzold et al., 2010) for quickly capturing and holding living zebrafish for rapid, high quality fluorescent expression in precise longitudinal imaging angles. The animals are easily inserted into a capillary of the correct diameter for the particular fish stage, then placed on the microscope and covered with a matched solution to remove the optical distortion from the capillary housing. The capillary is simply rotated for precise 0 degree (dorsal), 90 degree (lateral), and 180 degree (ventral) images on standard fluorescent microscopes, such as the Apotome. The technical bottleneck of standard microscopy to scan a whole embryo and larva is also labor-intensive and time-consuming. For instance, fluorescent scanning of a quarter of a 2dpf or 4dpf larvae required about 15 min exposure using a Zeiss Apotome microscope, one imaging modality deployed for this collection. To accelerate the expression profiling of these GBT lines, we subsequently utilized a Zeiss Lightsheet Z.1 SPIM microscope. The Lightsheet enabled high speed scanning of a whole embryo or larva, resulting in a nearly 20x faster image acquisition rate than the primary imaging process utilizing the Apotome. We prioritized and cataloged lines with robust expression at 2 and 4 days post fertilization (dpf). All zebrafish lines are freely available now through zfishbook (Clark et al., 2012) and are partially accessible from the Zebrafish International Resource Center. Consequently, we have generated 1138 lines by using each vector system showing in Supplemental Table 1), and updated results are posted at zfishbook.

Our throughput for cloning the GBT lines using traditional molecular methods was clearly an initial bottleneck. To help address this, we deployed a rapid cloning process based on methods used to isolate retroviral integrations (Varshney, Huang, et al., 2013; Varshney, Lu, et al., 2013). This method leverages the massive parallel sequencing technology of the Illumina MiSeq, yielding 101 bp sequencing reads, followed by a custom bioinformatics pipeline that involves both mapping and annotation. Fin-clips from four male animals per GBT locus are obtained during sperm cryopreservation and are used as a source of DNA. Shared inserts in multiple individuals from a single GBT line are considered candidate loci. This information is subsequently used to generate gene-specific primers to manually confirm linkage using 5‘ or 3’RACE, inverse PCR, or TAIL PCR to generate locus-specific primers for downstream molecular genotyping applications. 212 of the lines met the highest stringency of confirmed expression linkage and are classified as confirmed GBT integrations. An additional 143 lines have been initially tagged using this high throughput method, yielding candidate integration annotation as listed in zfishbook. The ability to complete the annotation status from candidate to confirmed for any given GBT locus with a desired expression profile is enhanced by the continued refinement of the zebrafish genome.

### mRFP Expression profiling reveals overlap with known annotation at 2dpf and substantive new expression data at both 2dpf and 4dpf

GBT lines are currently imaged to capture mRFP expression at both 2 and 4 dpf, including dorsal, sagittal and ventral views using the SCORE imaging method. (Petzold et al., 2010) In an openly accessible database of GBT lines, zfishbook, the images of mRFP expression pattern were stored within the media gallery associated with each line (Clark et al., 2012). We summarized published expression data of the tagged genes in both zfishbook and ZFIN in Table 1. Imaging the localization of transcripts and proteins at 4dpf is more difficult than those at 2 dpf, because accessibility of antisense RNA probes and antibodies into the larva’s body is technically limited for in the methods of both *in situ* hybridization and immunohistochemistry. Compared with the published data of gene expression in ZFIN, zfishbook currently provides almost the double number of genes with expression data at 2 dpf and 14 times the number of genes with expression data at 4 dpf (Table 1). In addition, zfishbook also provides novel expression data for 61 genes at any developmental stage (Fig.2).

**Figure 2.**
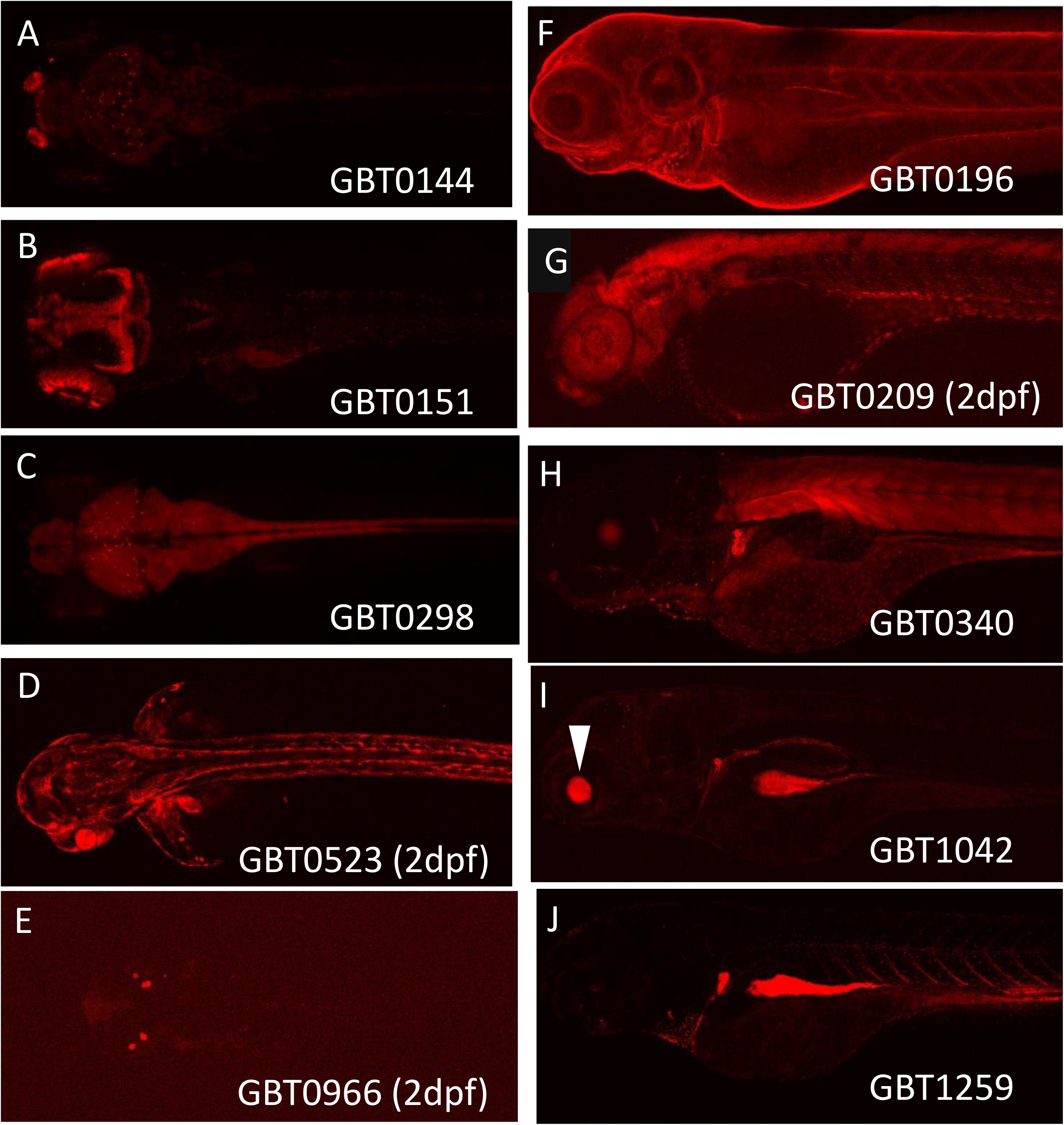
Novel protein expression. Representative novel expression data of trapped proteins fused to the mRFP reporter in this GBT collection. A. unkl localized in olfactory pit, cerebrum and spinal cord at 4dpf. B. nusap1 localized in retina and the top layer of both forebrain and midbrain at 4 dpf. C. zgc:194659 strongly expressed in the brain and spinal cord at 4 dpf. D. marcksl1a expressed in the lens, skin and notochord at 2 dpf. E. pipp2a specifically localized to otoliths at 2 dpf. F. ahnak specifically expressed in skin at 4 dpf. G. dph1 ubiquitously expressed showing granulized pattern in somites at 4 dpf. H. nfatc3a expressed in skeletal muscle and skin at 4 dpf. I. pard3bb localized to the pronephros and gut at 4 dpf. White arrowhead shows an artificial expression in lens driven by the promoter of 3’ exon trap of RP8.1. J. LOC100537272 expressed in circulatory cells in the blood stream at 4 dpf.

**Table 1.**
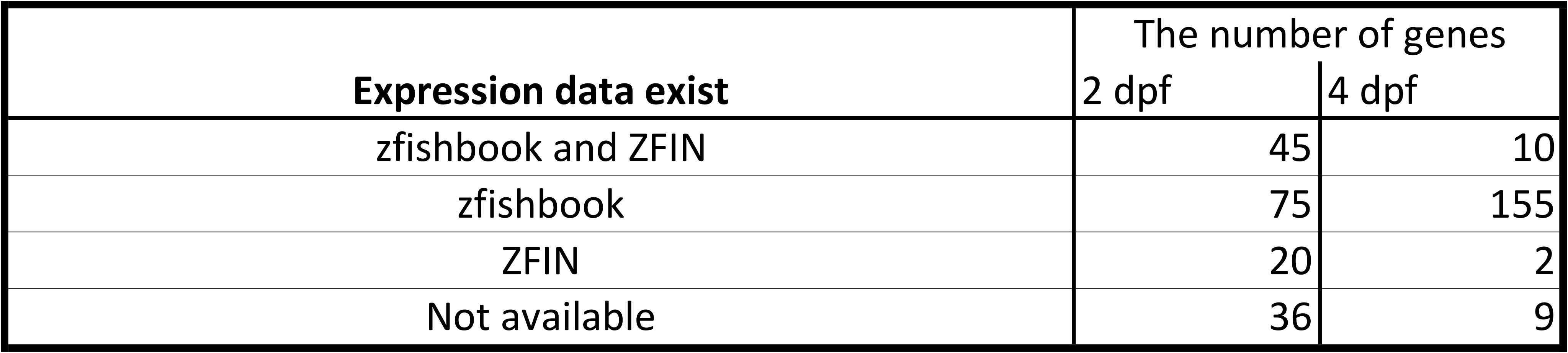
Comparison availability of expression data.

| Expression data exist | The number of genes |  |
| --- | --- | --- |
|  | 2 dpf | 4 dpf |
| zfishbook and ZFIN | 45 | 10 |
| zfishbook | 75 | 155 |
| ZFIN | 20 | 2 |
| Not available | 36 | 9 |

### High knockdown efficiency of endogenous transcripts induced by RP2

We directly compared published transposon insertional mutant vector systems (Fig. 3). The range and average knockdown levels in the FlipTrap system (Trinh le et al., 2011) produced a range of 4-30% (70-96% knockdown capacity) in six tested fish alleles, a similar range to our initial R-series protein trap vectors (R14-R15) that used a simple transcriptional terminator (Liao et al., 2012; Petzold et al., 2009) The pFT1 appears to be an improvement over these systems, in which the overall range and average read-through is reduced to 6-11% (89-94% knockdown) from four tested fish alleles (T. T. Ni et al., 2012). In contrast, the RP2.1 vector (Clark, Balciunas, et al., 2011), (Ding et al., 2013), (Ding et al., 2016) and this manuscript) maintains a strong knockdown (1% or less read-through) in 26 lines tested. Though deployed here using a nearly random, transposon-based delivery platform, the GBT vector system is an effective insertional mutagen suitable for an array of other – including targeted integration - genome-wide applications.

**Figure 3.**
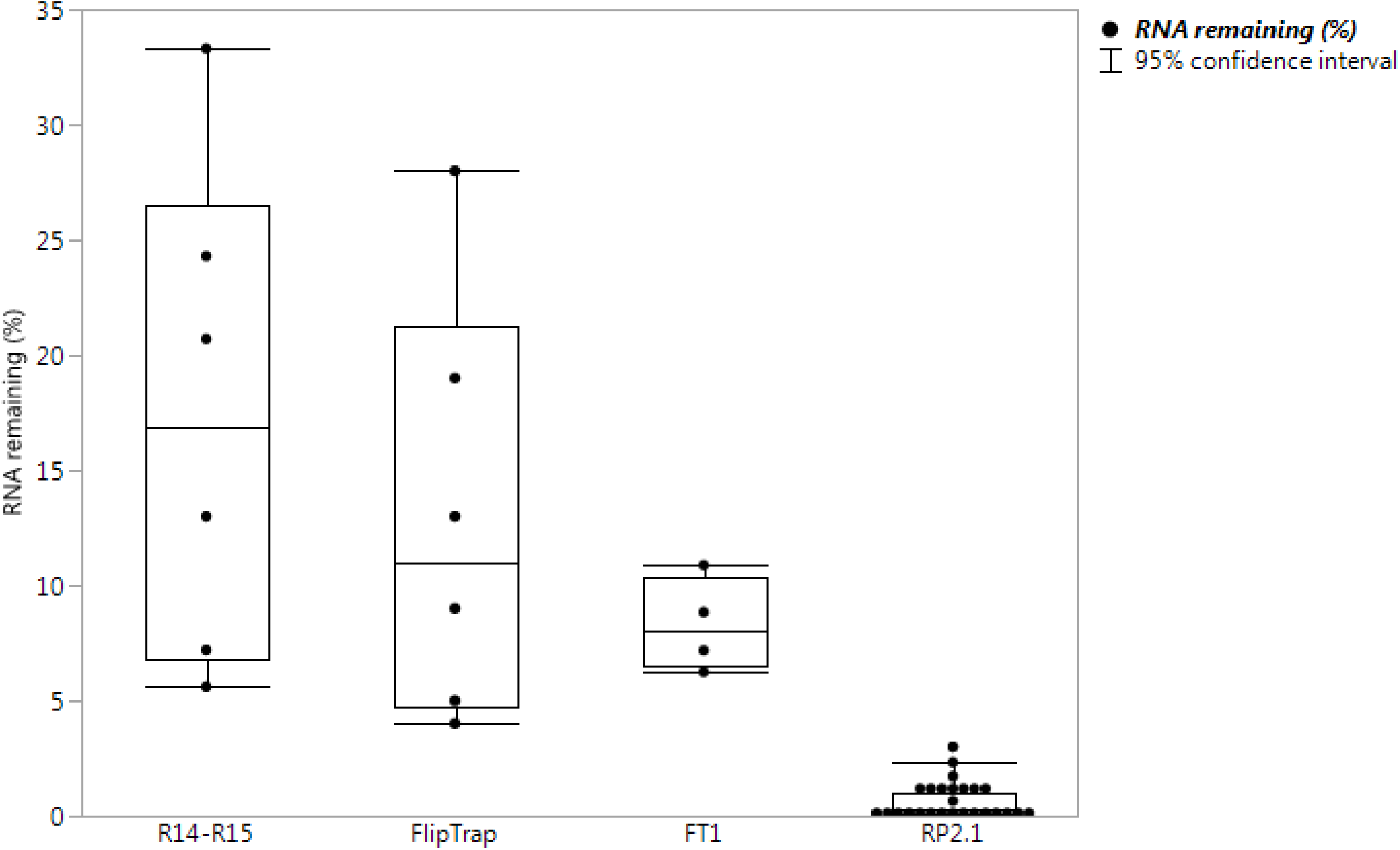
High knockdown efficiency of RP2.1 compared with other previous gene-trap systems. Black dots of bar graphs shows percentage of remaining endogenous transcripts in homozygous larvae with mean and 95% confidence interval indicated by individual lines. The data of previous protein trap systems were also converted from the data in the original articles, R14-R15, our initial R-series protein trap vectors (n= 6),(Clark, Balciunas, et al., 2011); FlipTrap, FlipTrap vectors (n= 6), (Trinh le et al., 2011); FT1, FT1 vector (n=4),(T. T. Ni et al., 2012); RP2.1 (n=26), (Clark, Balciunas, et al., 2011; Ding et al., 2013; El-Rass et al., 2017; Westcot et al., 2015) and unpublished data),

### Phenotypic Appearance Rate is Similar to Other Mutagenic Technologies for Forward Genetics Screening through 5 dpf

We conducted an initial forward genetic screen on embryos and early larvae of 179 RFP-positive GBT lines, identifying 12 recessive phenotypes, such as *ryr1b, fras1, tnnt2a, edar and hmcn1,*(Clark, Balciunas, et al., 2011; Westcot et al., 2015) visible during the first five days of development including lethality, heart, muscle, skin and other phenotypes. This 7% recovery of visible early developmental mutants is very similar to the 5% recovered visible mutants from the Sanger TILLING consortium analysis of truncated zebrafish genes (Kettleborough et al., 2013). This 7% is also comparable to prior retroviral (Amsterdam & Hopkins, 2004) and ENU(Haffter et al., 1996) zebrafish mutagenesis work that estimated between 1400 and 2400 genes (∼5-9% of the genome) would result in a visible embryonic phenotype when mutated.

### GBT alleles phenocopy known embryonic mutations

We tested the first five GBT lines in genes with known loss of function mutant phenotypes (*ryr1b*; *fras1*; *tnnt2a*; *edar; hmcn1*). All five of these alleles in genes with described loss of function defects are phenocopied by these GBT insertional alleles (Clark, Balciunas, et al., 2011); (Westcot et al., 2015); this manuscript). These loci represent a critical internal methods reference further validating the mutagenicity of these novel insertional vectors.

### Gene ontology analysis of GBT-tagged loci

To assess the diversity of GBT loci molecularly characterized to date, we utilized the PANTHER classification system (v.14.0, pantherdb.org, (Mi et al., 2013) and generated a table of protein class ontology tags in the molecularly isolated GBT lines. The PANTHER Protein Class ontology was adapted from the PANTHER INDEX (PANTHER/X) ontology that comprises two types of classifications: molecular function and biological process and includes commonly used classes of protein families. The molecular function schema classifies a protein based on its biochemical properties, such as receptor, cell adhesion molecule, or kinase. The biological process schema classifies a protein based on the cellular role or process in which it is involved, for example, carbohydrate metabolism (cellular role), TCA cycle (pathway), neuronal activities (process), or developmental processes (process) (Thomas PD et al, 2013). As of April 2018, almost half of known zebrafish genes (10626/25289 genes) were tagged in the PANTHER Protein Class ontology. 168 of our cloned GBT alleles mapped in the PANTHER system with 21 types of Protein Classes (Table 2). 18% and 16 % of the mapped GBT alleles are classified to nucleic acid binding (PC00171) and transcription factor (PC00218), respectively (Fig. 4). This result reveals that a quarter of the mapped genes possibly play a role in regulatory processes. Overall, however, the rich diversity of protein classes observed in our cloned traps suggests a large diversity will be represented by the overall collection and consistent with the random nature of genome integration events by the Tol2 transposon (Clark, Balciunas, et al., 2011).

**Figure 4.**
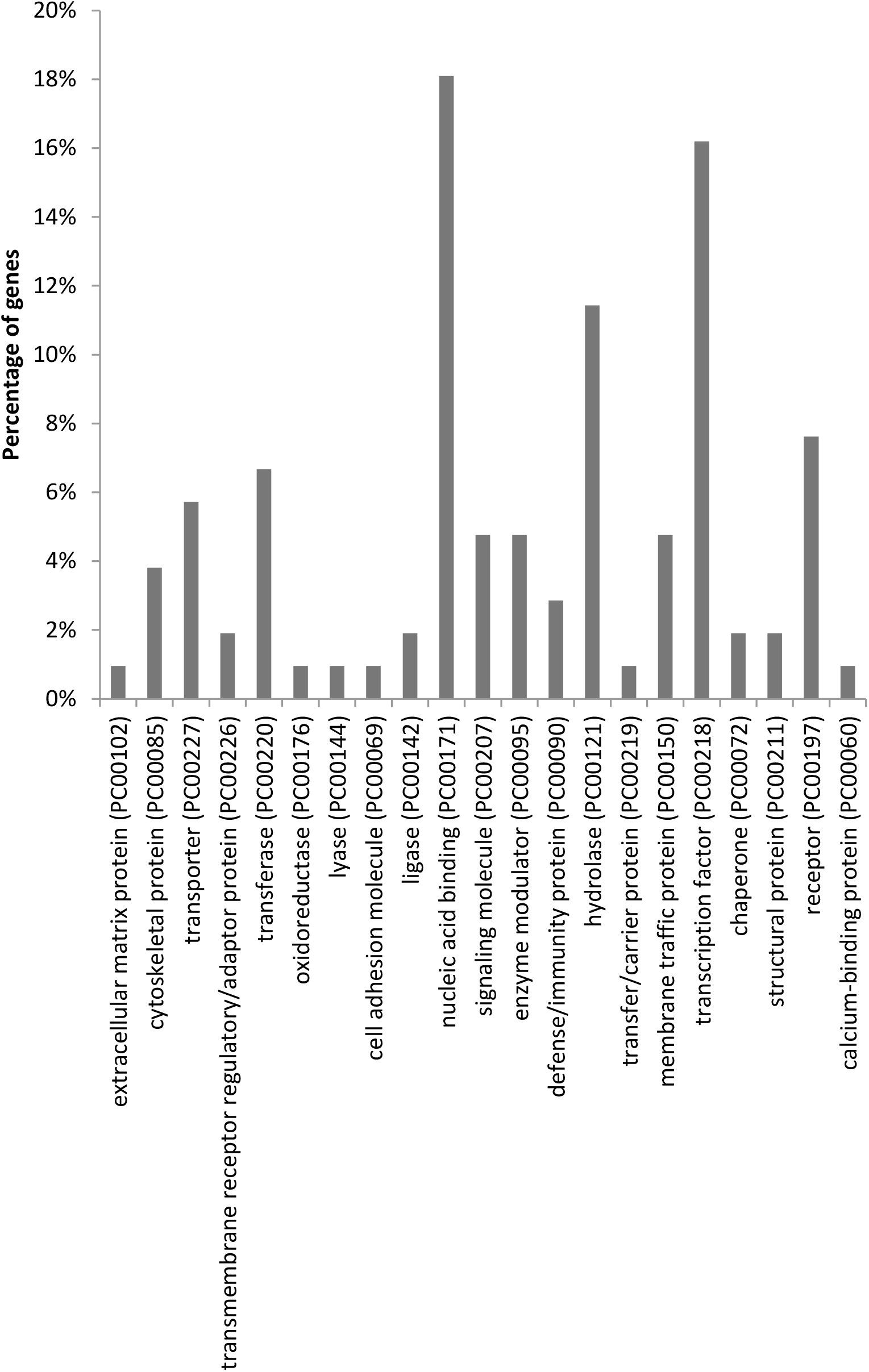
Summary of protein classes categorized the trapped proteins using PANTHER algorism. 77 trapped genes were successfully categorized at least one of 21 protein classes by using PANTHER gene ontology algorism. The details of the trapped genes classified in each protein class are listed in Table 2.

**Table 2.**
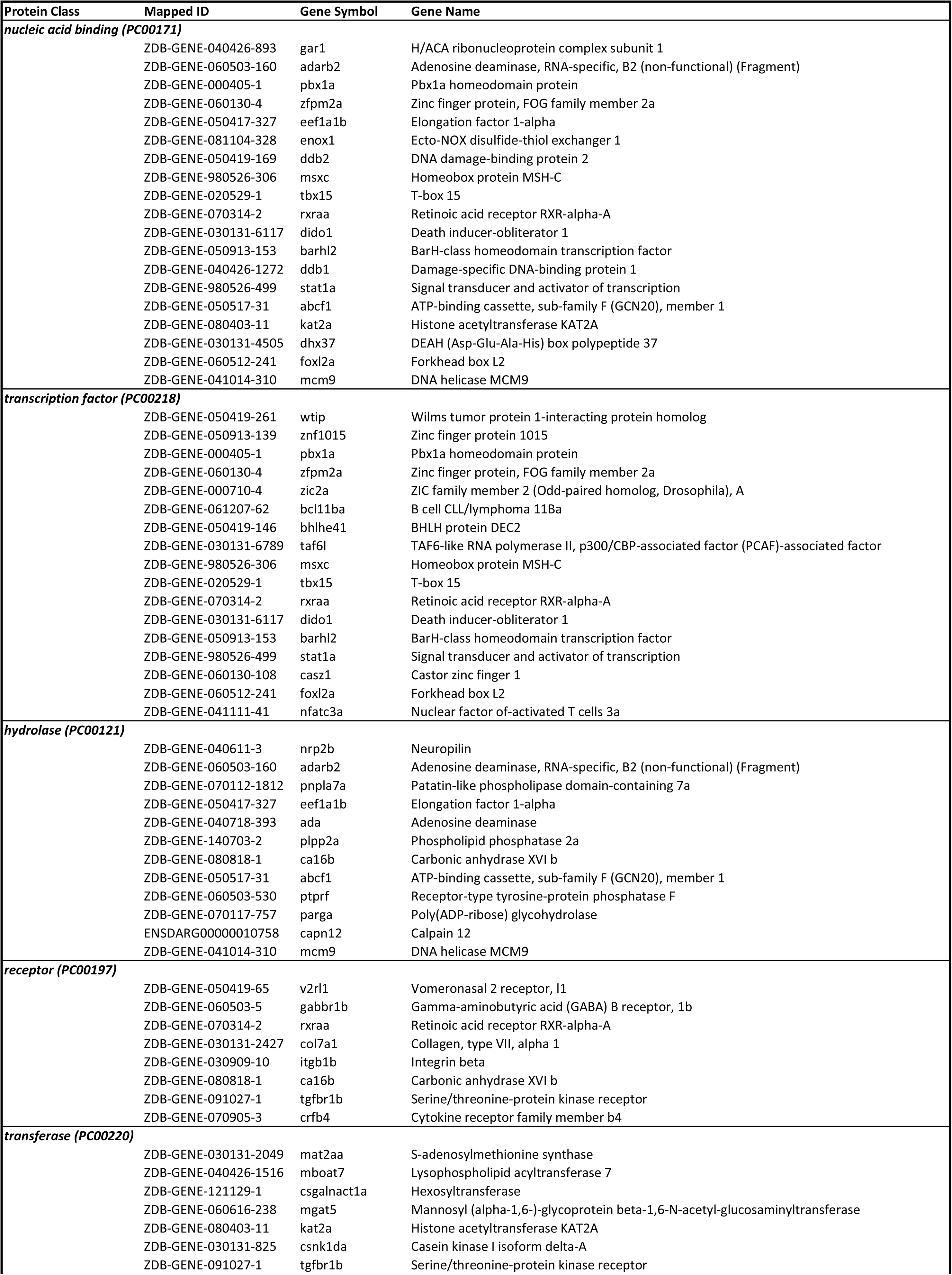
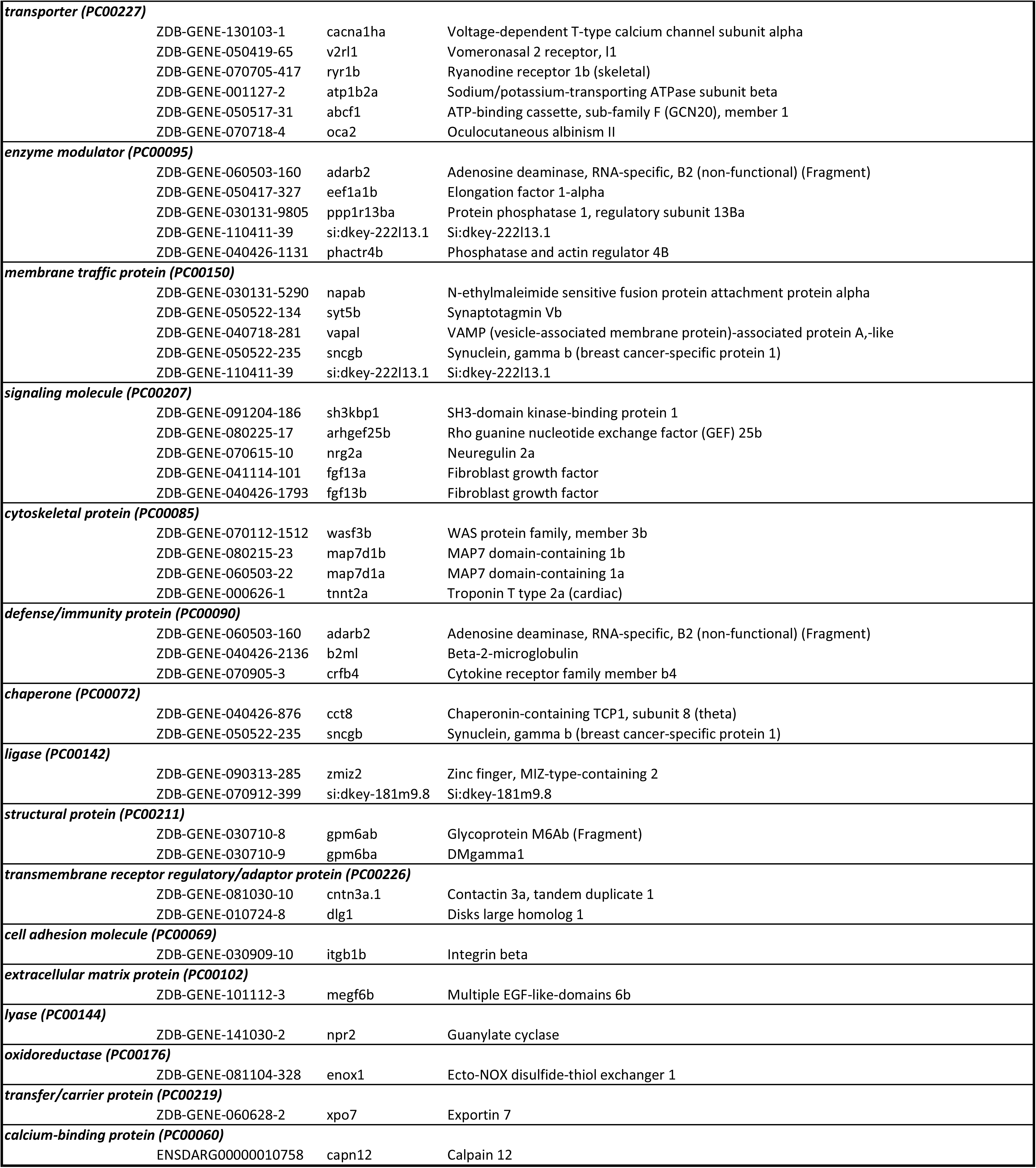
Protein classification.

### Disease-causing human orthologues of GBT-tagged genes

Of 183 GBT-tagged genes, 171 human orthologues were annotated in at least one public database such as ZFIN, Ensembl, Homologene, and InParanoid (Table 3). Several human orthologues were provisionally annotated using BLASTP and a synteny analysis tool, SynFind. In a previous study comparing the list of human genes possessing at least one zebrafish orthologue with the 3,176 genes bearing morbidity descriptions that are listed in the OMIM database, 82 % morbid genes 2,601 genes) can be related to at least one zebrafish orthologue (Howe et al., 2013). Surprisingly, 67 genes (about 37%) of 183 annotated human orthologues are associated with human disease involved in multi-organ system including nervous, circulatory, endocrine, metabolic, digestive, musculoskeletal, immune, and integument systems (Fig. 5 and Table 3) and many are not established in rodents and zebrafish (Table 5). The GBT protein-trap system provides a variety of potential human disease models which have a revertible allele that can interchange between disease and healthy cellular, organ and physiological states.

**Figure 5.**
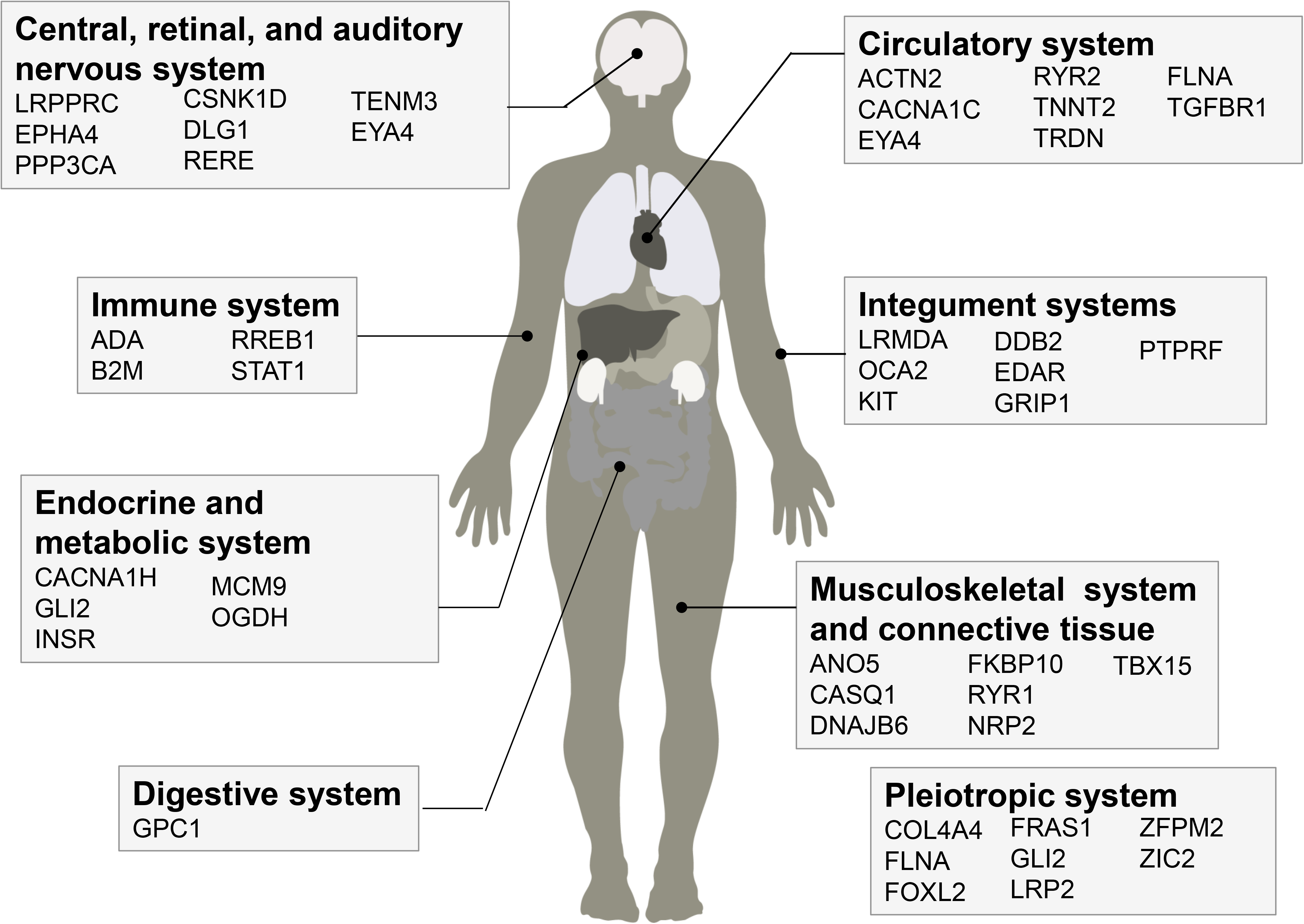
Disease-causing human orthologues of the trapped genes involved in human genetic disorders in multi-organ systems.

**Table 3.**
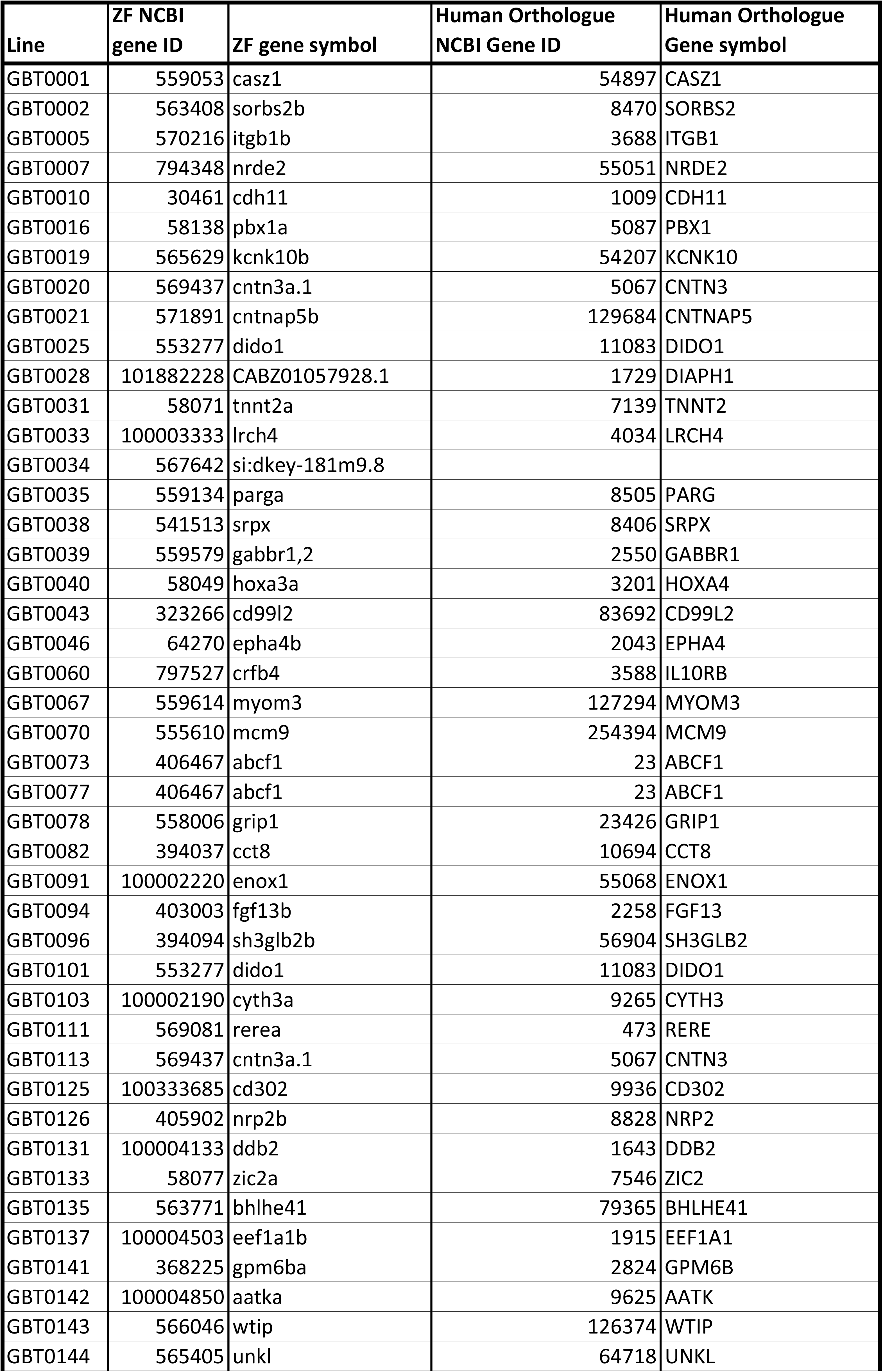
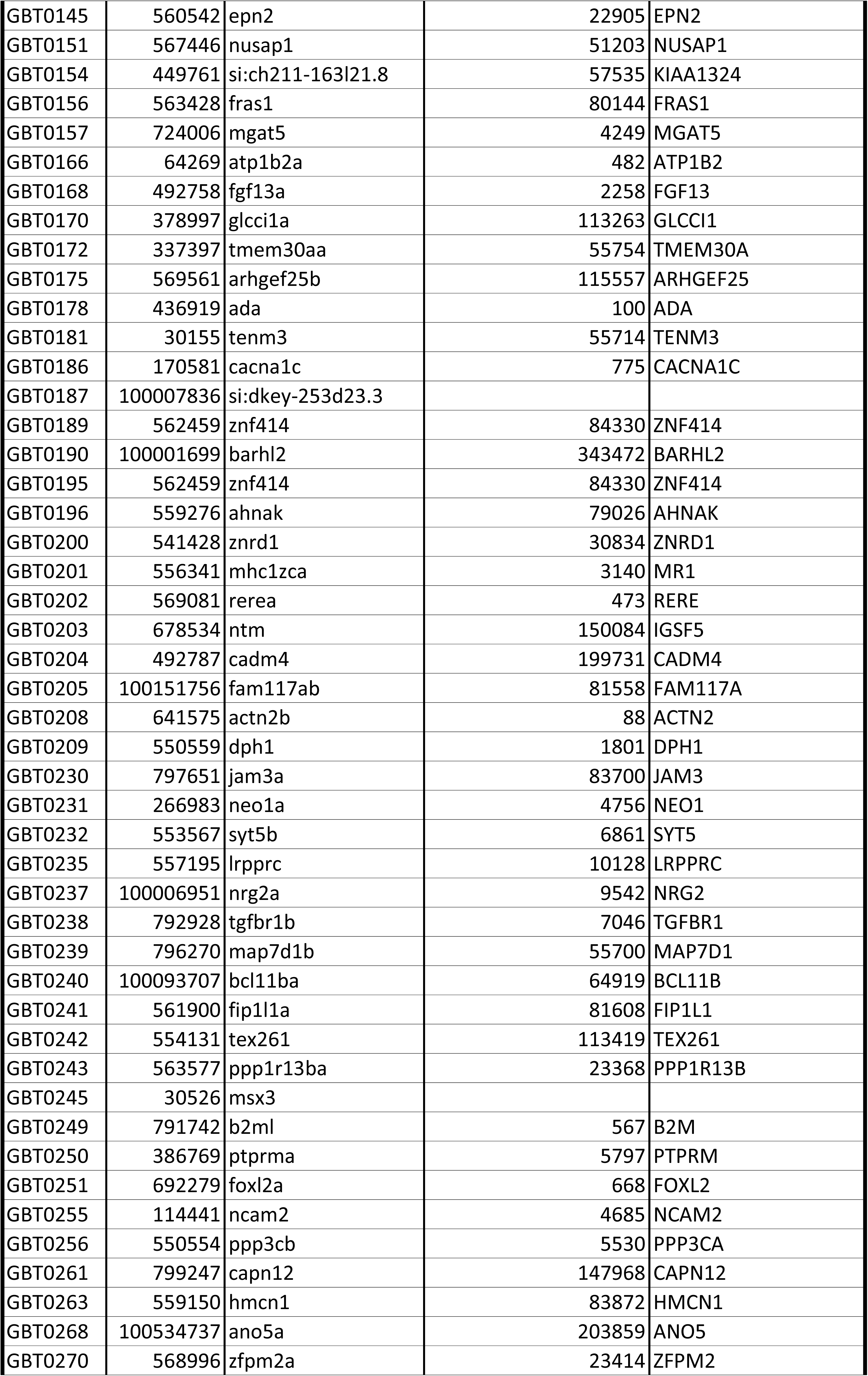
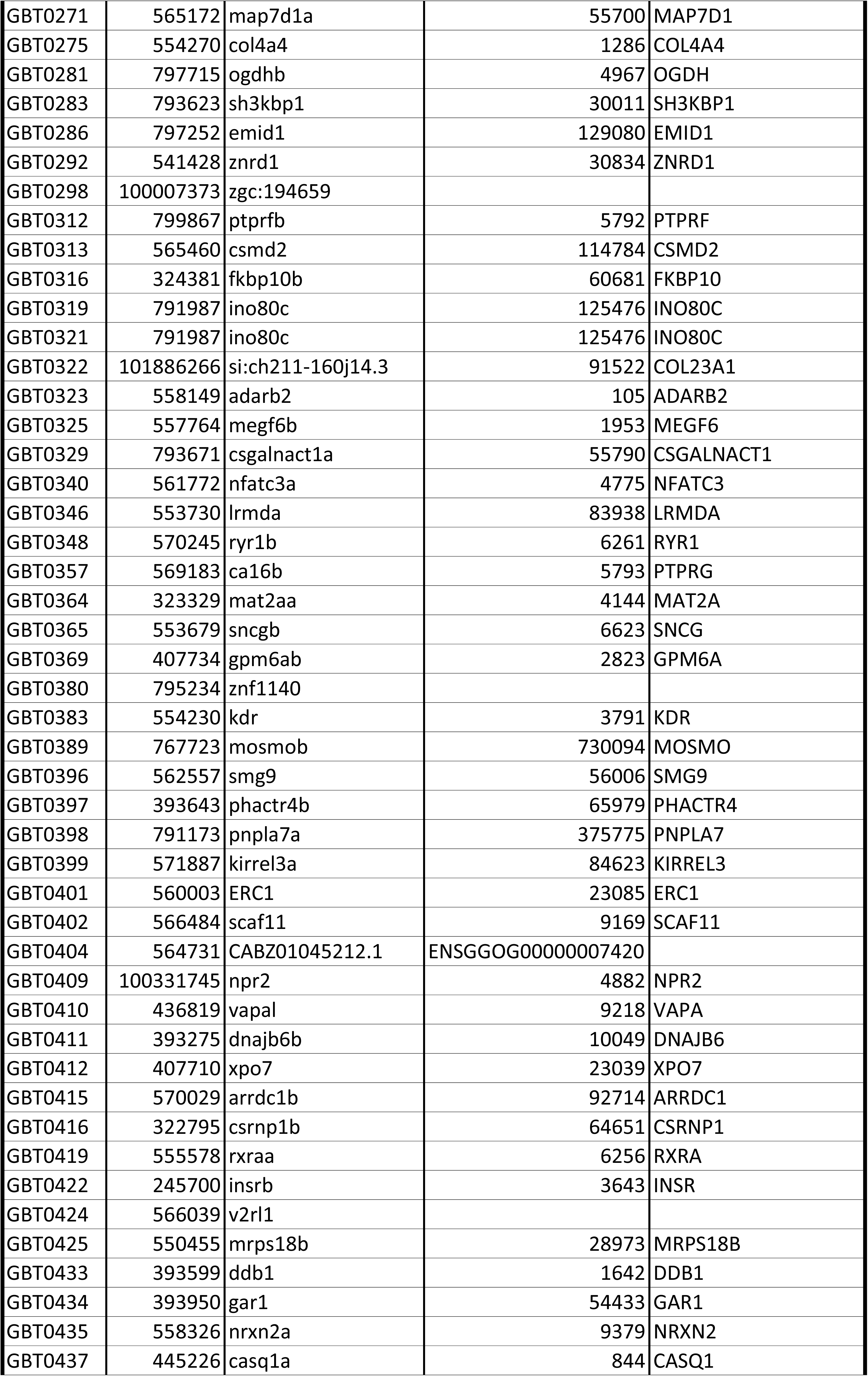
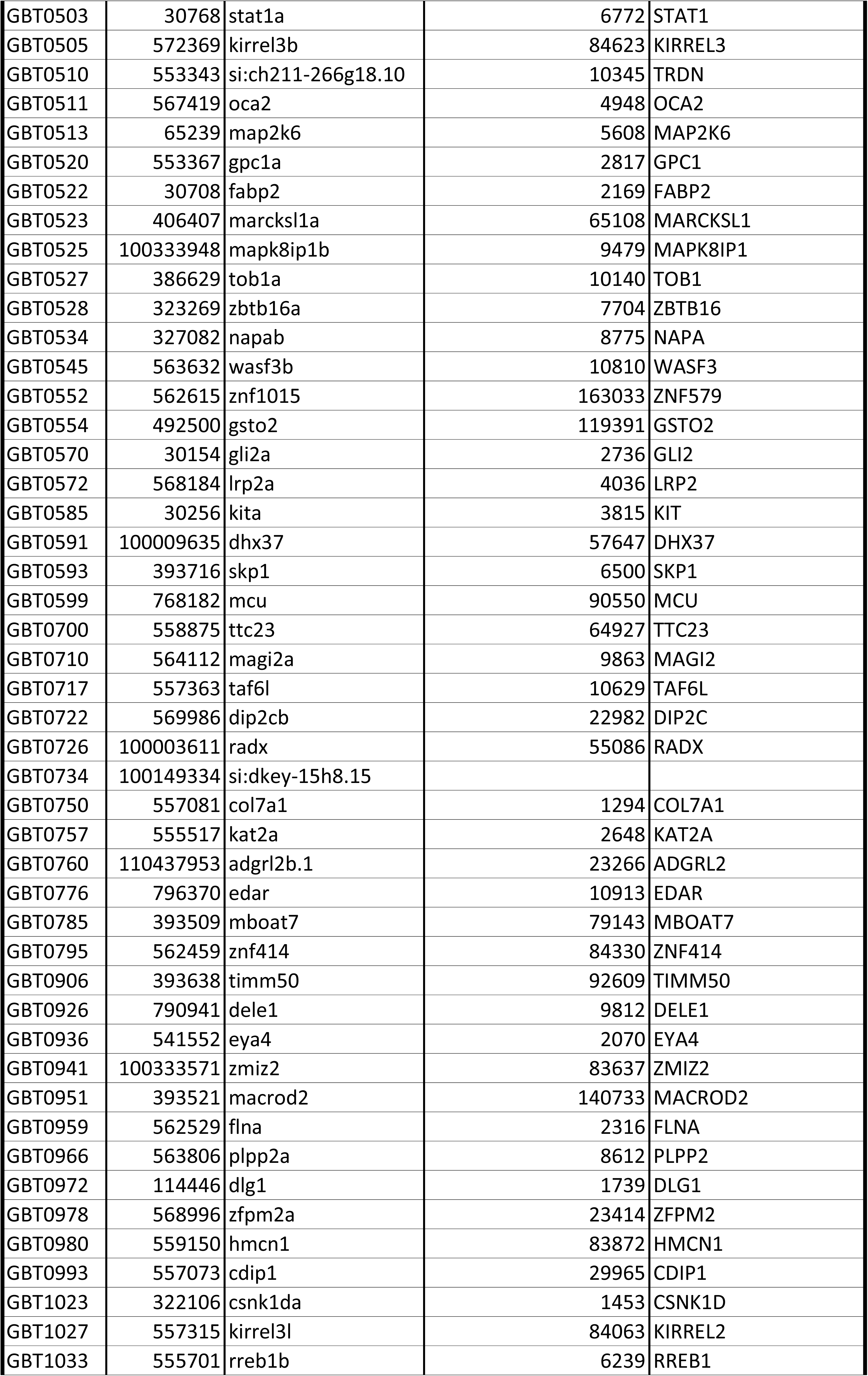
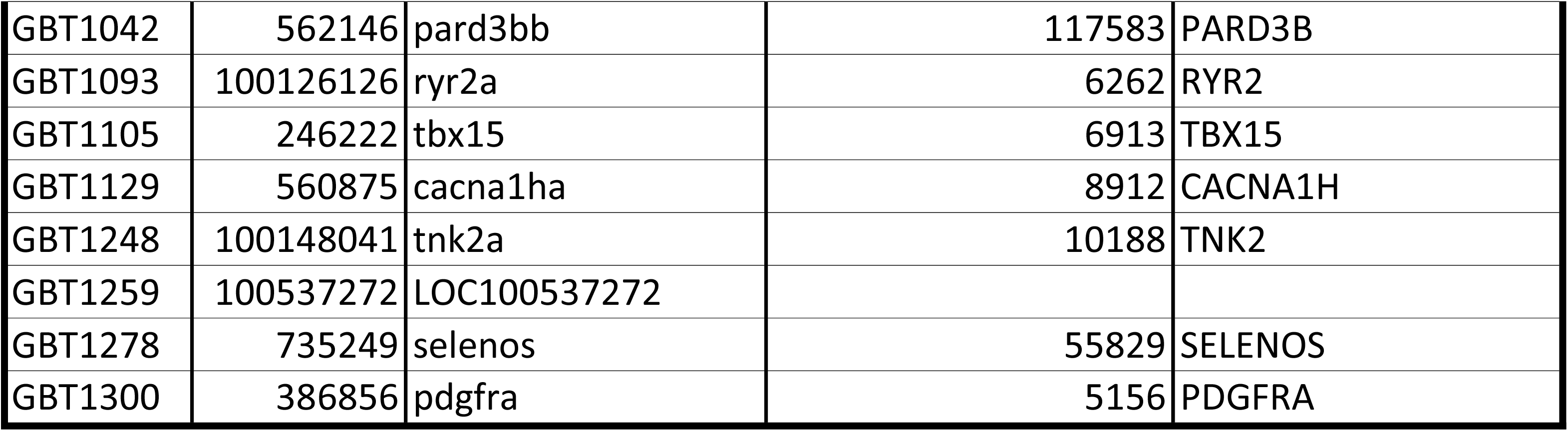
Disease-causing human orthologues.

### The GBT Protein Trap Reveals Protein-Coding Regions Not Predicted by the Zebrafish Genome Project

16 of 211 molecularly confirmed lines by manual, PCR-based mRFP linkage analysis using TALE, inverse, and RACE PCRs (Table 4) do not match any expressed sequence tag (EST) or predicted genes. However, in each case we were able to confirm transcription at the locus in wild-type animals, yielding new annotation for these loci in the zebrafish genome.

**Table 4.** Potential novel human disease models.

| GBT | Tagged Gene | Human Orthologue | Human Disease | Disease Models<br>(Number of models) | Reference<br>(PMID) |
| --- | --- | --- | --- | --- | --- |
| GBT0016 | pbx1a | PBX1 | CONGENITAL ANOMALIES OF KIDNEY AND URINARY TRACT SYNDROME WITH ORWITHOUT HEARING LOSS<br>ABNORMAL EARS OR DEVELOPMENTAL DELAY | mouse (1) | 12591246 |
| GBT0031 | tnnt2a | TNNT2 | CARDIOMYOPATHY DILATED 1D | mouse (5) | 18606313,<br>27936050,<br>17556660,<br>18349139 |
| GBT0031 | tnnt2a | TNNT2 | CARDIOMYOPATHY FAMILIAL HYPERTROPHIC 2 | mouse (9) | 16326803,<br>9788962,<br>11171784,<br>18349139,<br>10449439,<br>9637714,<br>23532597 |
| GBT0078 | grip1 | GRIP1 | FRASER SYNDROME 3 | mouse (2) | 14730302,<br>16880404 |
| GBT0131 | ddb2 | DDB2 | XERODERMA PIGMENTOSUM COMPLEMENTATION GROUP E | mouse (4) | 14769931,<br>15558025 |
| GBT0133 | zic2a | ZIC2 | HOLOPROSENCEPHALY 5 | mouse (3) | 27466203,<br>18617531,<br>10677508 |
| GBT0135 | bhlhe41 | BHLHE41 | SHORT SLEEPER | mouse (1) | 19679812 |
| GBT0156 | fras1 | FRAS1 | FRASER SYNDROME 1 | mouse (5) | 12766769,<br>12766770,<br>24143185,<br>15623520,<br>15838507 |
| GBT0178 | ada | ADA | SEVERE COMBINED IMMUNODEFICIENCY AUTOSOMAL RECESSIVE TCELL-NEGATIVE B CELL-NEGATIVE NK CELL-<br>NEGATIVE DUE TO ADENOSINEDEAMINASE DEFICIENCY | mouse (1) | 9478961 |
| GBT0186 | cacna1c | CACNA1C | TIMOTHY SYNDROME | mouse (1) | 21878566 |
| GBT0240 | bcl11ba | BCL11B | IMMUNODEFICIENCY 49 | zebrafish (1) |  |
| GBT0251 | foxl2a | FOXL2 | Blepharophimosis ptosis and epicanthus inversus | mouse (3) | 15056605,<br>14736745,<br>24565867 |
| GBT0268 | ano5a | ANO5 | MUSCULAR DYSTROPHY LIMB-GIRDLE AUTOSOMAL RECESSIVE 12 | mouse (1) | 26911675 |
| GBT0270 | zfpm2a | ZFPM2 | Tetralogy of Fallot | mouse (1) | 10892744,<br>12223418 |
| GBT0270 | zfpm2a | ZFPM2 | DIAPHRAGMATIC HERNIA 3 | mouse (1) | 16103912 |
| GBT0275 | col4a4 | COL4A4 | ALPORT SYNDROME AUTOSOMAL RECESSIVE | mouse (4) | 21196518,<br>24522496 |
| GBT0348 | ryr1b | RYR1 | CENTRAL CORE DISEASE OF MUSCLE | mouse (3) | 25564733,<br>19959667,<br>7515481 |
| GBT0396 | smg9 | SMG9 | HEART AND BRAIN MALFORMATION SYNDROME | mouse (1) | 27018474 |
| GBT0409 | npr2 | NPR2 | Acromesomelic dysplasia Maroteaux type | mouse (2) | 23065701,<br>17728275 |
| GBT0411 | dnajb6b | DNAJB6 | autosomal dominant limb-girdle muscular dystrophy type 1 | zebrafish (1-NOT), mouse (1) | 26362252 |
| GBT0422 | insrb | INSR | DONOHUE SYNDROME | mouse (2-NOT) |  |
| GBT0511 | oca2 | OCA2 | ALBINISM OCULOCUTANEOUS TYPE II | rabbit (1) | Magnussen,<br>K. (1952). |
| GBT0572 | lrp2a | LRP2 | DONNAI-BARROW SYNDROME | mouse (1) | 20653565 |
| GBT0585 | kita | KIT | MASTOCYTOSIS CUTANEOUS | mouse (5) | 24788138,<br>21148330 |
| GBT0585 | kita | KIT | Piebald trait | mouse (1) | 20095975 |
| GBT0585 | kita | KIT | Gastrointestinal stromal tumor | mouse (5) | 16061643,<br>22652566,<br>18098338,<br>12754375 |
| GBT0710 | magi2a | MAGI2 | NEPHROTIC SYNDROME TYPE 15 | mouse (1) | 25271328 |
| GBT0750 | col7a1 | COL7A1 | recessive dystrophic epidermolysis bullosa | mouse (4) | 18382769,<br>19893033,<br>10523500 |
| GBT0776 | edar | EDAR | hypohidrotic ectodermal dysplasia | mouse (1) | 10431242,<br>17148670,<br>9799835 |
| GBT0776 | edar | EDAR | HAIR MORPHOLOGY 1 | mouse (1) | 18561327 |
| GBT0785 | mboat7 | MBOAT7 | MENTAL RETARDATION AUTOSOMAL RECESSIVE 57 | mouse (1) | 23097495 |
| GBT0959 | flna | FLNA | Periventricular nodular heterotopia 1 | mouse (1) | 16825286 |
| GBT1023 | csnk1da | CSNK1D | ADVANCED SLEEP PHASE SYNDROME FAMILIAL 2 | mouse (2) | 15800623 |
| GBT1093 | ryr2a | RYR2 | Ventricular tachycardia catecholaminergic polymorphic 1 with or without atrial dysfunction and/or dilated<br>cardiomyopathy | mouse (10) | 27482086,<br>25775566,<br>26121139,<br>24755079,<br>23152493,<br>22828895,<br>20224043,<br>18419777,<br>16873551,<br>15890976 |
| GBT1093 | ryr2a | RYR2 | ARRHYTHMOGENIC RIGHT VENTRICULAR DYSPLASIA FAMILIAL 2 | mouse (1) | 16873551 |
| GBT1105 | tbx15 | TBX15 | COUSIN SYNDROME | mouse (1) | 19068278 |

**Table 5.** Novel transcripts.

| Line | Vector | Genome location | Genome version |
| --- | --- | --- | --- |
| GBT0115 | RP2 | chr8:12923125-12923133 | ZV9 |
| GBT0129 | RP2 | chr12:45451589-45451597 | ZV9 |
| GBT0148 | RP2 | chr16:55188302-55188310 | ZV9 |
| GBT0264 | RP2 | chr5:49960612-49960620 | ZV9 |
| GBT0506 | RP2 | chr18:46186350-46186358 | ZV9 |
| GBT0573 | RP2 | chr6:33630487-33630495 | GRCz10 |
| GBT0586 | RP2 | chr13:9112806-9112814 | GRCz10 |
| GBT0702 | RP2 | chr13:9112806-9112814 | GRCz10 |
| GBT0724 | RP8 | chr7:7633071-7633079 | GRCz10 |
| GBT0994 | RP2 | chr3:39627361-39627369 | GRCz10 |
| GBT1024 | RP2 | chr25:5175242-5175250 | GRCz10 |
| GBT1071 | RP8 | chr9:21818261-21818269 | GRCz10 |
| GBT1100 | RP2 | chr5:25509243-25509251 | GRCz10 |
| GBT1116 | RP2 | chr14:14724098-14724106 | GRCz10 |
| GBT1168 | RP2 | chr5:39664405-39664412 | GRCz10 |

## Discussions

### Novel Expression Annotation Revealed By Protein Trapping of Endogenous Genes

The RP2 and RP8 vectors of GBT system were assembled to capture all three reading frames of fusion protein of the trapped gene with the mRFP reporter. This GBT system reveals that mRFP-truncated fusion proteins exhibit distinct subcellular localization. This is particularly noteworthy for the 4dpf stages because published gene expression data at late developmental stages have been limited by the technical difficulties in conducting such analyses using traditional techniques such as whole mount *in situ* hybridization (WISH) at these larval and later time points. We note that the mRFP-truncated fusion protein may localize ectopically in cases where the protein localization signal is contained in the C-terminal domain (Clark, Balciunas, et al., 2011; Trinh le & Fraser, 2013). Although the extent that subcellular localization recapitulates the endogenous protein is dependent on each insertion locus specifics, the visualizing and illuminating spatiotemporal expression patterns of trapped protein may facilitate dynamic studies of specific cell types and molecular functions in a living vertebrate. The novel expression description at 4dpf in nearly five of six cloned loci demonstrates the dearth of 3D expression annotation for the overwhelming number of genes, even in one of the most studied model system such as the zebrafish. With ever-improving microscope-based imaging tools (Liu et al., 2018), these lines have the potential to help annotate at diverse developmental and adult stages, while also potentially imaging subcellular expression for a subset of protein trap fusions.

### Gene-Break Protein Trap in an Effective Insertional Mutagen with High Knockdown Efficiency and Cre-Reversion to WT allele

Transposons offer several unique features over and above traditional static mutational approaches, including high quality expression tools and new regulated mutagenesis methodologies. From the perspective of genome engineering development, GBT technology was the first method for revertible allele generation of vertebrates outside of the mouse (Clark, Balciunas, et al., 2011); (Ding et al., 2013). We know two major potential biases that may yield non-random trapping coverage of the genome. First, the RP2.1 protein trap was initially designed around a single reading frame. Upon molecular analysis of our first lines, however, we discovered that RP2.1 encodes a second, alternative splice acceptor yielding protein trap expression from a second reading frame due to this alternative splicing event in a significant number of our lines. The deliberate development of RP2 and RP8 vectors for each reading frame obviates this potential limitation when used in other delivery modalities besides the Tol2 transposon. For example, these vectors would be suitable for gene editing-based targeted knockin methods.

The GBT system deployed here has been joined by two new and complementary transposon mutagenesis systems. The FlipTrap system by Dr. Fraser’s group(Trinh le et al., 2011), which can be mutagenic when provided Cre recombinase, is primarily focused on imaging fusion proteins in vivo and addressing cellular dynamics and related questions. The FT1 system by the Chen lab is a complementary flipping trap that can use either Cre or Flp recombinases to regulate alleles depend on the original orientation of the insertion (T. T. Ni et al., 2012). GBT-based zebrafish alleles are highly complementary and non-redundant to those generated by other mutagenesis methods, including these other transposons and demonstrated higher knockdown efficiency of the WT transcripts than those mutagens because of the use of an enhanced polyadenylation signal and a putative boundary element between 5’ protein trap and 3’ exon trap cassettes. Importantly, since the initiation of this project to generate a collection of Cre-revertible mutant alleles, several groups have now reported collections of tissue-specific Cre driver lines including the Brand lab (Jungke et al., 2013; http://crezoo.crt-dresden.de/crezoo/), the Zcre consortium (http://zcre.org.uk/) and the Wen lab in the PTC Consortium. The GBT system described here is a two-component, molecularly regulatable mutagenesis approach that offers the ability to test for the sufficiency of protein-encoding loci in regulated, tissue- and cell-specific applications.

### Functional Diversity of the Trapped Proteins by the GBT system

To analyze distribution of protein functions of the trapped genes by GBT system, we performed GO analysis using the PANTHER protein classification. Although the protein functions related in transcriptional regulatory process, such as nucleic acid binding and transcription factors represented one relatively common class of isolated genes, the protein GO analysis indicated that the GBT protein trap was a useful tool for capturing a wide range of protein functions in addition to cell fate regulators and related nuclear genes.

### Cloned GBT Loci Represent a Rich Collection of Potential Human Disease Models

More than 7,000 human diseases have already been described, and 80% of those are thought to have a genetic origin (Varga et al., 2018). Model organism studies can be a pivotal resource for understanding gene function which possibly provides additional insight into the cause of particular disease, thereby contributing to understanding of the pathogenic process and discovery of the therapeutic strategy (Wangler et al., 2017). Annotation of human orthologues of 160 tagged genes revealed that GBT technology yield a high frequency potential human disease models with loss of gene function, tracking expression of the truncated protein and Cre-revertible mutated allele to rescue the phenotype.

Furthermore, the modeling human genetic diseases using the GBT system has advantages compared with reverse genetic approaches such as TALEN and CRISPR-Cas9 systems. In initial phenotype screening, PR2.1 mutagenesis demonstrated 7% phenotype appearance as much as the other ENU- and retroviral forward genetic screenings (Amsterdam & Hopkins, 2004; Haffter et al., 1996; Kettleborough et al., 2013). For example, the annotation of disease-causing human orthologues of the tagged genes also revealed that the GBT system comprehensively developed mutants in zebrafish orthologues of human disease loci, including nervous, cardiovascular, endocrine, digestive, musculoskeletal, immune, and integument systems. Surprisingly, this system generated 68 of pioneering mutants in orthologs of human disease loci (Supplemental Table 2).

### Discovery of Novel Transcripts by Trapping Unpredicted Genes

Since the completion of the zebrafish reference genome sequencing, it has enabled many new discoveries to be made, in particular the positional cloning of hundreds genes from mutation affecting embryogenesis behavior, physiology, and health and disease. However, a few poorly assembled regions remain (Howe et al., 2013). In molecular cloning of GBT lines generated, we found that a surprising proportion of the sequenced insertions does not correspond to any predicted genes. Although we have not formally excluded that mRFP expression might, in some case, be an artifact, the data of gene prediction provided in genome databases reveals some prediction errors. These results suggest that the algorithms used to predict genes from genome databases have missed a significant number of genes. The protein trapping by using GBT system may useful in identifying unsuspected novel genes, expressions and functions *in vivo* in real time.

### New Genomic Insights Using the GBT Random Insertion Mutagen

The ready ease of these mRFP-based protein traps for basic expression analyses demonstrates how much we do not know about our overall proteome and the codex that is our genome. Nearly forty percent and five of six of the cloned genes show no expression at 2dpf and 4dpf, respectively, in ZFIN. At the subcellular level, such protein traps in conjunction with new microscopy techniques represent just another method for cellular and mechanistic analyses in an *in vivo* cellular context. Although these were made using random insertional approaches, new targeted integration tools using gene editing such as GeneWeld(Wierson et al., 2018) should readily empower labs to build their custom GBT lines in the future for genes not in this collection. Together, this initial 1100+ GBT collection is a new contribution to the use of the zebrafish to annotate the vertebrate genome.

## Author contributions

SCE and KJC conceived research; NI, SCE, KJC for basic experimental design including detailed analyses of the collection; MS, CD, MU, SER, YD, RU, MM, JJE, XX, DB, SCE and KJC generated GBT collection, NI, KS, MS, LG, CD, MU, RU, YD, SAF, WL and KJC conducted phenotype screening of GBT mutant lines; NI, KS, MS, LG, CD, MU, RU, YD, WL and KJC conducted molecular biology analyses; GV and SB conducted next generation sequencing; NI, KS, SCE and KJC conducted bioinformatics-based analyses; LAS, NI, KJC and SCE conducted comparative genomics analyses, NI, KS, MS, LG, KJS and SCE wrote the manuscript; SCE, KJC, JJE, XX, MH, SAF, XYW, SB, XX and SL consulted to this research.

## Funding

Supported by NIH grants (GM63904, DA14546, DK093399 and HG 006431), Natural Sciences and Engineering Research Council of Canada, grant RGPIN 05389-14 and the Mayo Foundation.

## Competing interests

The authors declare no competing interests.

## Acknowledgements

This work is supported by grants from the National Institutes of Health (GM63904; DA14546; HG006431) and the Mayo Foundation. We thank Zoltan Varga for sharing of the ZIRC sperm cryopreservation protocol prior to publication. Appreciation is also extended to the Mayo Clinic Zebrafish Facility staff for their excellent support. This research was in part funded by the Intramural Research Program of the National Human Genome Research Institute; National Institutes of Health (S.M.B.: 1ZIAHG000183)

---

Supplemental Table 1 Generation of 11 hundred independent lines by all three reading frames of mRFP reporter in both RP2 and RP8 cassettes.

Supplemental Table 2 Disease-causing human orthologues of the trapped protein Supplemental Figure 1 Representative expression patterns of mRFP fusion protein by RP2 and RP8 integration

