## Supplementary figures and images for "The Vertebrate Codex Gene Breaking Protein Trap Library For Genomic Discovery and Disease Modeling Applications"

### Supplemental Figure 1

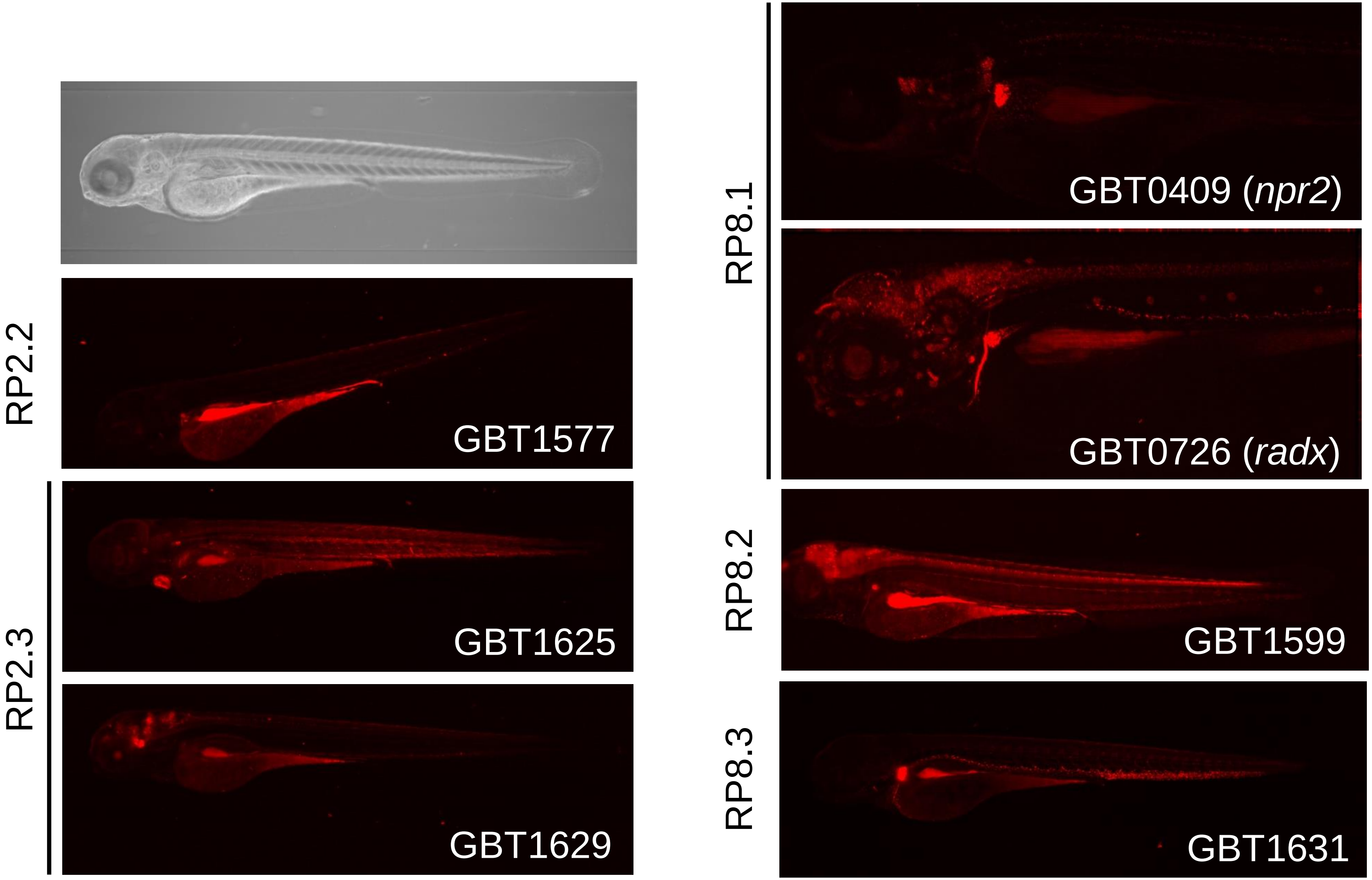


Supplemental Figure 1
