## Supplemental Table1 for "The Vertebrate Codex Gene Breaking Protein Trap Library For Genomic Discovery and Disease Modeling Applications"

| **Vector** | **Number of lines** |
|---|---|
| R14/ R14.5 | 7 |
| R15 | 18 |
| R16 | 9 |
| RP2.1/ RP2 | 905 |
| RP2.2 | 7 |
| RP2.3 | 30 |
| RP8.1/RP8 | 128 |
| RP8.2 | 20 |
| RP8.3 | 14 |
| Total | 1138 |

### Supplemental Table 1
