## Supplemental Table 2 for "The Vertebrate Codex Gene Breaking Protein Trap Library For Genomic Discovery and Disease Modeling Applications"

| Gene name | Organ system | Phenotype description | Study reference of phenotype | Source name of phenotype |
|---|---|---|---|---|
| **ACTN2** | Circulatory system | CARDIOMYOPATHY DILATED 1AA WITH OR WITHOUT LEFT VENTRICULAR NONCOMPACTION | 612158 | OMIM |
| **ADA** | Immune system | SEVERE COMBINED IMMUNODEFICIENCY AUTOSOMAL RECESSIVE TCELL-NEGATIVE B CELL-NEGATIVE NK CELL-NEGATIVE DUE TO ADENOSINEDEAMINASE DEFICIENCY | 102700 | OMIM |
| **ANO5** | Musculoskeletal system and connective tissue | MIYOSHI MUSCULAR DYSTROPHY 3 | 613319 | OMIM |
| **ANO5** | Musculoskeletal system and connective tissue | MUSCULAR DYSTROPHY LIMB-GIRDLE AUTOSOMAL RECESSIVE 12 | 611307 | OMIM |
| **ANO5** | Musculoskeletal system and connective tissue | GNATHODIAPHYSEAL DYSPLASIA | 166260 | OMIM |
| **B2M** | Immune system | IMMUNODEFICIENCY 43 | 241600 | OMIM |
| **CACNA1C** | Circulatory system | BRUGADA SYNDROME 3 | 611875 | OMIM |
| **CACNA1C** | Circulatory system | TIMOTHY SYNDROME | 601005 | OMIM |
| **CACNA1H** | Endcrine and metabolic system | HYPERALDOSTERONISM FAMILIAL TYPE IV | 617027 | OMIM |
| **CASQ1** | Musculoskeletal system and connective tissue | Myopathy vacuolar with casq1 aggregates | 616231 | OMIM |
| **COL4A4** | Pleiotropic system | ALPORT SYNDROME AUTOSOMAL RECESSIVE | 203780 | OMIM |
| **CSNK1D** | Central retinal and auditory nervous system | ADVANCED SLEEP PHASE SYNDROME FAMILIAL 2 | 615224 | OMIM |
| **DDB2** | Integument system | XERODERMA PIGMENTOSUM COMPLEMENTATION GROUP E | 278740 | OMIM |
| **DLG1** | Central retinal and auditory nervous system | Cleft lip/palate | 199306 | Orphanet |
| **DNAJB6** | Musculoskeletal system and connective tissue | MUSCULAR DYSTROPHY LIMB-GIRDLE AUTOSOMAL DOMINANT 1 | 603511 | OMIM |
| **EDAR** | Integument system | ECTODERMAL DYSPLASIA 10B HYPOHIDROTIC/HAIR/TOOTH TYPE AUTOSOMALRECESSIVE | 224900 | OMIM |
| **EDAR** | Integument system | ECTODERMAL DYSPLASIA 10A HYPOHIDROTIC/HAIR/NAIL TYPE AUTOSOMAL DOMINANT | 129490 | OMIM |
| **ERC1** | Central retinal and auditory nervous system | Distal monosomy 12p | 280325 | Orphanet |
| **EYA4** | Central retinal and auditory nervous system | DEAFNESS, AUTOSOMAL DOMINANT 10 | 601316 | OMIM |
| **EYA4** | Circulatory system | CARDIOMYOPATHY, DILATED, 1J | 603550 | OMIM |

| | | | | |
|---|---|---|---|---|
| **FKBP10** | Musculoskeletal system and connective tissue | OSTEOGENESIS IMPERFECTA TYPE XI | 610968 | OMIM |
| **FKBP10** | Musculoskeletal system and connective tissue | Bruck syndrome 1 | 259450 | OMIM |
| **FLNA** | Pleiotropic system | Periventricular nodular heterotopia 1 | 300049 | OMIM |
| **FLNA** | Circulatory system | Cardiac valvular dysplasia X-linked | 314400 | OMIM |
| **FLNA** | Pleiotropic system | Intestinal pseudoobstruction neuronal chronic idiopathic X-linked | 300048 | OMIM |
| **FLNA** | Pleiotropic system | MELNICK-NEEDLES SYNDROME | 309350 | OMIM |
| **FLNA** | Pleiotropic system | FRONTOMETAPHYSEAL DYSPLASIA 1 | 305620 | OMIM |
| **FLNA** | Pleiotropic system | OTOPALATODIGITAL SYNDROME TYPE I | 311300 | OMIM |
| **FLNA** | Pleiotropic system | OTOPALATODIGITAL SYNDROME TYPE II | 304120 | OMIM |
| **FLNA** | Pleiotropic system | TERMINAL OSSEOUS DYSPLASIA | 300244 | OMIM |
| **FOXL2** | Pleiotropic system | Blepharophimosis ptosis and epicanthus inversus | 110100 | OMIM |
| **FRAS1** | Pleiotropic system | FRASER SYNDROME 1 | 219000 | OMIM |
| **GLI2** | Pleiotropic system | HOLOPROSENCEPHALY 9 | 610829 | OMIM |
| **GLI2** | Pleiotropic system | CULLER-JONES SYNDROME | 615849 | OMIM |
| **GLI2** | Endcrine and metabolic system | Combined pituitary hormone deficiencies genetic forms | 95494 | Orphanet |
| **GPC1** | Digestive system | Isolated Biliary atresia | 30391 | Orphanet |
| **GRIP1** | Integument system | FRASER SYNDROME 3 | 617667 | OMIM |
| **GRIP1** | Integument system | FRASER SYNDROME 1 | 219000 | OMIM |
| **INSR** | Endcrine and metabolic system | DIABETES MELLITUS INSULIN-RESISTANT WITH ACANTHOSIS NIGRICANS | 610549 | OMIM |
| **INSR** | Endcrine and metabolic system | HYPERINSULINEMIC HYPOGLYCEMIA FAMILIAL 5 | 609968 | OMIM |
| **INSR** | Endcrine and metabolic system | PINEAL HYPERPLASIA INSULIN-RESISTANT DIABETES MELLITUS AND SOMATIC ABNORMALITIES | 262190 | OMIM |
| **INSR** | Endcrine and metabolic system | DONOHUE SYNDROME | 246200 | OMIM |
| **KIT** | Integument system | MASTOCYTOSIS CUTANEOUS | 154800 | OMIM |
| **LRMDA** | Integument system | ALBINISM OCULOCUTANEOUS TYPE VII | 615179 | OMIM |
| **LRP2** | Pleiotropic system | DONNAI-BARROW SYNDROME | 222448 | OMIM |

| | | | | |
|---|---|---|---|---|
| **LRPPRC** | Central retinal and auditory nervous system | LEIGH SYNDROME FRENCH CANADIAN TYPE | 220111 | OMIM |
| **MCM9** | Endcrine and metabolic system | OVARIAN DYSGENESIS 4 | 616185 | OMIM |
| **NPR2** | Musculoskeletal system and connective tissue | Acromesomelic dysplasia Maroteaux type | 602875 | OMIM |
| **OCA2** | Integument system | ALBINISM OCULOCUTANEOUS TYPE II | 203200 | OMIM |
| **OGDH** | Endcrine and metabolic system | ALPHA-KETOGLUTARATE DEHYDROGENASE DEFICIENCY | 203740 | OMIM |
| **PPP3CA** | Central retinal and auditory nervous system | Undetermined early-onset epileptic encephalopathy | 442835 | Orphanet |
| **PTPRF** | Integument system | BREASTS AND/OR NIPPLES APLASIA OR HYPOPLASIA OF 2 | 616001 | OMIM |
| **RERE** | Central retinal and auditory nervous system | NEURODEVELOPMENTAL DISORDER WITH OR WITHOUT ANOMALIES OF THE BRAIN EYE OR HEART | 616975 | OMIM |
| **RREB1** | Immune system | 22q11.2 Deletion Syndrome | 567 | Orphanet |
| **RYR1** | Musculoskeletal system and connective tissue | MINICORE MYOPATHY WITH EXTERNAL OPHTHALMOPLEGIA | 255320 | OMIM |
| **RYR1** | Musculoskeletal system and connective tissue | MYOPATHY CONGENITAL WITH FIBER-TYPE DISPROPORTION | 255310 | OMIM |
| **RYR1** | Musculoskeletal system and connective tissue | CENTRAL CORE DISEASE OF MUSCLE | 117000 | OMIM |
| **RYR2** | Circulatory system | Ventricular tachycardia catecholaminergic polymorphic 1 with or without atrial dysfunction and/or dilated cardiomyopathy | 604772 | OMIM |
| **RYR2** | Circulatory system | ARRHYTHMOGENIC RIGHT VENTRICULAR DYSPLASIA FAMILIAL 2 | 600996 | OMIM |
| **STAT1** | Immune system | IMMUNODEFICIENCY 31C | 614162 | OMIM |
| **STAT1** | Immune system | IMMUNODEFICIENCY 31A | 614892 | OMIM |
| **STAT1** | Immune system | IMMUNODEFICIENCY 31B | 613796 | OMIM |
| **TBX15** | Musculoskeletal system and connective tissue | COUSIN SYNDROME | 260660 | OMIM |
| **TENM3** | Central retinal and auditory nervous system | MICROPHTHALMIA ISOLATED WITH COLOBOMA 9 | 615145 | OMIM |
| **TGFBR1** | Circulatory system | LOEYS-DIETZ SYNDROME 1 | 609192 | OMIM |
| **TNNT2** | Circulatory system | CARDIOMYOPATHY DILATED 1D | 601494 | OMIM |
| **TNNT2** | Circulatory system | CARDIOMYOPATHY FAMILIAL RESTRICTIVE 3 | 612422 | OMIM |

| | | | | |
|---|---|---|---|---|
| **TRDN** | Circulatory system | Ventricular tachycardia catecholaminergic polymorphic 1 with or without atrial dysfunction and/or dilated cardiomyopathy | 604772 | OMIM |
| **ZFPM2** | Pleiotropic system | Tetralogy of Fallot | 187500 | OMIM |
| **ZFPM2** | Pleiotropic system | 46XY sex reversal 9 | 616067 | OMIM |
| **ZFPM2** | Pleiotropic system | DIAPHRAGMATIC HERNIA 3 | 610187 | OMIM |
| **ZIC2** | Pleiotropic system | HOLOPROSENCEPHALY 5 | 609637 | OMIM |

Supplemental Table 2
